# PNMA2 forms non-enveloped virus-like capsids that trigger paraneoplastic neurological syndrome

**DOI:** 10.1101/2023.02.09.527862

**Authors:** Junjie Xu, Simon Erlendsson, Manvendra Singh, Matthew Regier, Iosune Ibiricu, Gregory S. Day, Amanda L. Piquet, Stacey L. Clardy, Cedric Feschotte, John A. G. Briggs, Jason D. Shepherd

## Abstract

The *paraneoplastic Ma antigen* (*PNMA*) genes are associated with cancer-induced paraneoplastic syndromes that present with neurological symptoms and autoantibody production. How PNMA proteins trigger a severe autoimmune disease is unclear. *PNMA* genes are predominately expressed in the central nervous system with little known functions but are ectopically expressed in some tumors. Here, we show that *PNMA2* is derived from a Ty3 retrotransposon that encodes a protein which forms virus-like capsids released from cells as non-enveloped particles. Recombinant PNMA2 capsids injected into mice induce a robust autoimmune reaction with significant generation of autoantibodies that preferentially bind external “spike” PNMA2 capsid epitopes, while capsid-assembly-defective PNMA2 protein is not immunogenic. PNMA2 autoantibodies present in cerebrospinal fluid of patients with anti-Ma2 paraneoplastic neurologic disease show similar preferential binding to PNMA2 “spike” capsid epitopes. These observations suggest that PNMA2 capsids released from tumors trigger an autoimmune response that underlies Ma2 paraneoplastic neurological syndrome.

## Introduction

The *PNMA* family of genes are predominately expressed in the brain (*1-4*), but little is known about their biological function. *PNMA1-2* were first identified as genes encoding proteins that are targets of neuronal autoantibodies in blood and cerebrospinal fluid (CSF) from patients with certain paraneoplastic neurological syndromes (*1-3*). Patients with elevated levels of PNMA1 (Ma1) or PNMA2 (Ma2) autoantibodies experience limbic/brainstem encephalitis and cerebellar degeneration (*5*). These patients often present with solid peripheral tumors (*1, 6, 7*) that are a potential source of PNMA protein (*8-11*) and antibodies to PNMA proteins can be diagnostic of specific cancers (*12*). Prognosis in this paraneoplastic syndrome is poor (*7*). It has remained enigmatic why these proteins elicit such a strong autoimmune response, but given the severity of disease, it is imperative to gain an understanding of the pathophysiology to better target immunomodulatory treatments.

The PNMA family of genes is predicted to have Gag-homology domains (*13, 14*). Recent studies identified Gag-derived proteins co-opted from long terminal repeat (LTR) retrotransposons that mediate a new and unexpected virus-like intercellular communication pathway (*13-17*). These genes include the neuronal *Arc* genes that are critical for synaptic plasticity, memory, and cognition (*15, 16, 18, 19*). The Gag domain contained in LTR retrotransposons and retroviruses is necessary and sufficient to produce viral-like particles that package their mRNA during replication. The *Arc* gene, which was co-opted from a Ty3/mdg4 (formerly known as Ty3/gypsy) retrotransposon in the common ancestor of tetrapod vertebrates, encodes a protein that has retained virus-like biology. Arc assembles into virus-like capsids that are released from neurons in membrane-enveloped extracellular vesicles (EVs) that transfer RNA and protein cell-to-cell (*16*). The Drosophila Arc (dArc) homologs also assemble capsids from a Gag domain that are released in EVs (*16, 20*), but originated independently from a distinct lineage of Ty3/mdg4 retrotransposons (*18, 21*). The Gag-like gene *PEG10*, which was derived from yet another lineage of Ty3/mdg4 retrotransposons during mammalian evolution, has also retained ancestral virus-like properties such as RNA binding, capsid formation, and release in EVs (*14, 22-25*). Together these studies suggest that a diverse set of co-opted Gag genes have preserved biochemical properties of ancestral retroelements (*26*), but their physiological functions and role in human diseases are poorly characterized.

In this study, we investigated whether putative virus-like properties of PNMA proteins trigger an autoimmune reaction that underlies paraneoplastic neurological disorders. Concentrating on the *PNMA2* gene, we find that *PNMA2* is normally exclusively and highly expressed in neurons. PNMA2 protein self-assembles into virus-like capsids that are released from cells as non-enveloped capsids. Based on these results, we posited that PNMA2 capsids released outside the central nervous system (CNS) may cause autoantibody generation. Consistent with this, mice injected with PNMA2 capsids produce strong autoantibody generation without adjuvant. Based on atomic resolution structures of PNMA2 capsids, we designed PNMA2 mutants that are unable to form capsids. Injection of capsid mutant protein does not elicit autoantibody production. Finally, we find that PNMA2 autoantibodies from paraneoplastic patients preferentially bind to exterior surface “spike” epitopes of PNMA2 capsids. Taken together, our results show that PNMA2 is a Gag-like protein that forms endogenous virus-like capsids that are released without a membrane, which can trigger a robust immune response when ectopically expressed outside the CNS. These properties may underlie the neurological deficits associated with Ma2 paraneoplastic disease.

## Results

### PNMA2 evolved from a Ty3/mdg4 retrotransposon coopted in the ancestor of placental mammals

To trace the evolutionary origins of *PNMA2*, we conducted phylogenomic analysis. The *PNMA2* gene is conserved at an orthologous genomic position across all major lineages of placental mammals but absent from the genomes of marsupials and non-mammalian species (Figure 1a). Thus, *PNMA2* was coopted in the common ancestor of placental mammals ∼100 million years ago. The *PNMA2* gene is composed of three exons. The first two are short noncoding exons interrupted by long introns, while the third exon encodes the open reading frame corresponding to the Gag-derived portion of an ancient MamGyp-int element of the Ty3/mdg4 superfamily (*16, 21*). The predicted *PNMA2* promoter region is located only ∼150 bp away from its nearest neighboring gene, DPYSL2, which is arranged in the opposite orientation relative to *PNMA2* (Figure 1a). This arrangement suggests that *PNMA2* and *DPYSL2* share a bidirectional promoter. Because the promoter region and the *DPYSL2* gene are more deeply conserved across vertebrate species than *PNMA2* (see sequence conservation track in Figure 1a), it is likely that the cooption of *PNMA2* was facilitated by the capture of a bidirectional promoter from a pre-existing neighboring gene, *DPYSL2*.

**Figure 1.**
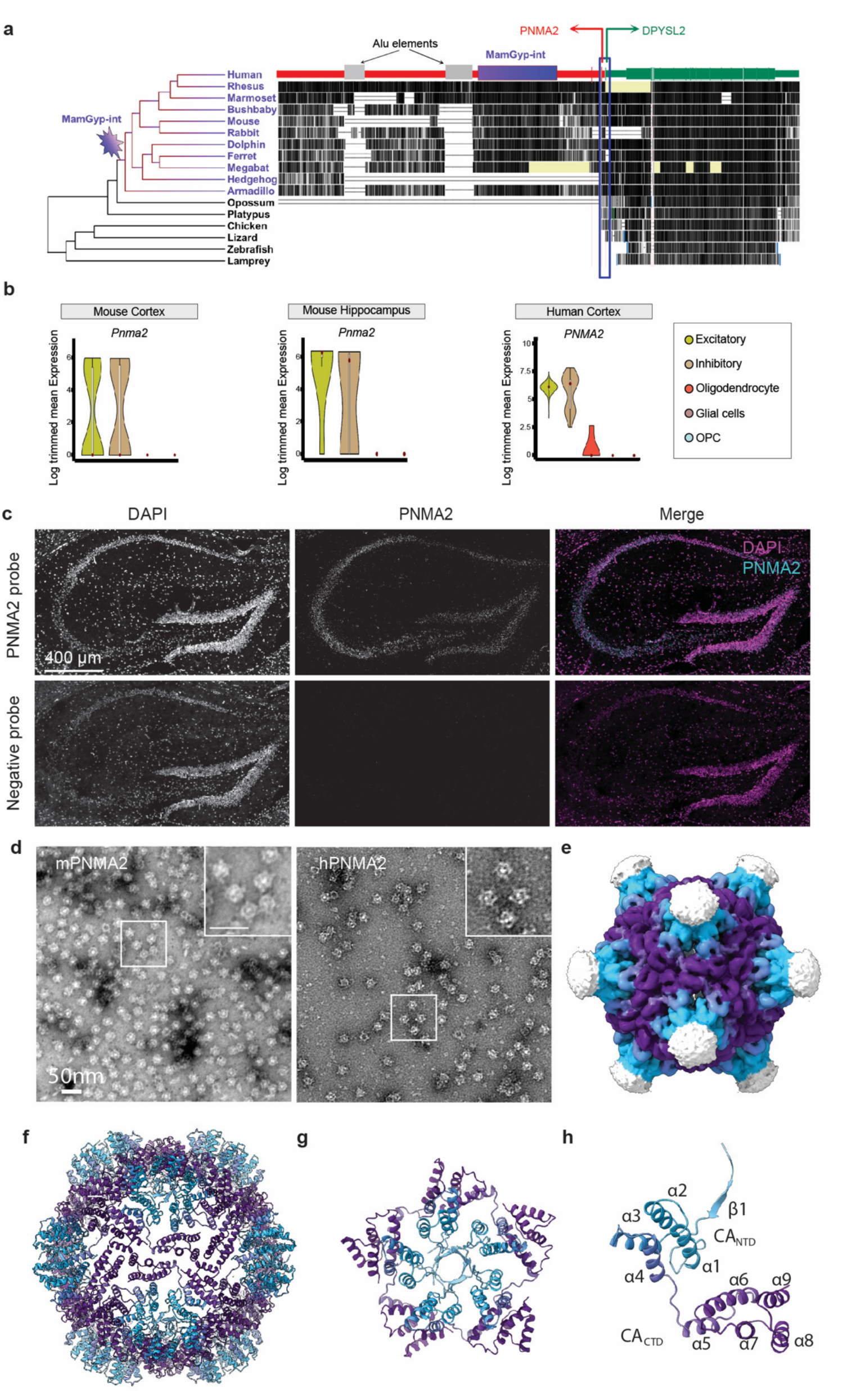
Ty3/mdg4 retrotransposon-derived *PNMA2* encodes proteins self-assembling into virus-like capsids. **a**. Structure and evolution of PNMA2. The figure is derived from the Vertebrate Multiz Alignment and Conservation track at the UCSC Genome Browser (hg38) and shows the phylogenetic relationship (left) and level of sequence conservation (right) for a subset of vertebrate genomes across the mRNA sequence and shared promoter region of *PNMA2* (red) and *DPYSL2* (green) gene models. There are three transposable elements annotated by RepeatMasker within the *PNMA2* mRNA: a Mam-Gyp-int element (purple box) that gave rise to the Gag-like coding sequence, and two Alu elements (grey boxes) embedded within the 3’ UTR. The arrows depict the predicted transcription start sites for *PNMA2* and *DPYSL2*. The conservation track shows that *PNMA2* mRNA sequence is deeply conserved across placental mammals (except for the two Alu elements which are primate-specific insertions), but not other vertebrates, while *DPYSL2* and promoter region are conserved more deeply across vertebrate evolution pointing at their earlier origins. **b**. Single-cell expression of *PNMA2* mRNA in the human cortex (right panel) and mouse cortex and hippocampus (left panel). **c**. PNMA2 single molecule fluorescent *in situ* hybridization probe (RNAscope) and negative control probe were used to detect PNMA2 mRNA in wild-type mouse (2 months old) hippocampal slice. Scale bar: 400µm. **d**. Representative negative-stained EM images of purified recombinant mouse PNMA2 (mPNMA2) capsids and human PNMA2 (hPNMA2) capsids. Scale bar: 50nm. **e**. Surface representation of mPNMA2 as resolved from cryo-EM, viewed down the two-fold axis. The spike densities are not resolved. **f**. Atomic model of the T=1 mPNMA2 capsid. CAµis depicted in purple and the CAµin cyan. **g**. External view of the isolated five-fold capsomere. **h**. One of 60 mPNMA2 monomers required to form the T=1 capsids.

### PNMA2 is expressed in mammalian neurons

To gain insight into *PNMA2* expression and localization, we first surveyed bulk RNA-seq data from the GTEx project (*27*), which reveals that *PNMA2* is highly and almost exclusively expressed in human brain tissue samples (gtexportal.org/home/gene/ENSG00000240694) (Supplementary Figure S1). Next, we analyzed single-cell RNA-sequencing datasets to examine *PNMA2* expression in human motor cortex, mouse cerebral cortex, and mouse hippocampus (*28-31*). This analysis reveals that *PNMA2* mRNA is expressed in excitatory and inhibitory neurons, with very low expression in glia and oligodendrocyte lineages (Fig 1b). To confirm the RNA-seq data, we used fluorescent *in situ* hybridization (RNAScope) and found that *PNMA2* mRNA is highly transcribed in mouse brain (Figure 1c) and primary cultured hippocampal neurons (Supplementary Figure S2). To determine whether *PNMA2* expression pattern is conserved across primates, we analyzed single-cell RNA-seq data from brain samples representing 33 anatomical regions from humans (N=132), chimpanzees (N=96), and macaques (N=96)) (*32*). *PNMA2* expression is remarkably conserved in grey matter areas of all three species, suggesting that the expression of *PNMA2* in the brain is under strong purifying selection (Supplemental Figure S3).

### PNMA2 proteins self-assemble into virus-like capsids

Purified recombinant mammalian and Drosophila Arc proteins spontaneously form virus-like capsids (*15, 16*). PNMA 3, 5, and 6a also form virus-like particles (*14*). To determine if PNMA2 also forms capsids, we purified recombinant PNMA2 in an *E. coli* expression system and imaged the samples by negative stain EM. Purified mouse PNMA2 (mPNMA2) and human PNMA2 (hPNMA2) protein spontaneously form ordered virus-like capsids (Figure 1d). PNMA2 capsids were highly stable, even when treated with high concentrations of detergent (Supplemental Figure S4). Using single-particle cryo-EM we reconstructed density maps of the entire icosahedral mPNMA2 capsid as well as of local regions of the capsid surface after symmetry relaxation at a resolution of 3.2 Å (Figure 1e, Supplemental Figure S5-6). mPNMA2 forms icosahedral capsids with an outer diameter of ∼200 Å and a triangulation number T = 1, composed of 12 pentameric capsomeres and 60 individual mPNMA2 molecules (Figure 1f). The protein shell of the capsid is roughly 30 Å thick. Based on the obtained maps we built an atomic model of the mPNMA2 capsid (Figure 1f; Supplemental Figure S7; Supplementary Movie 1; Supplementary Table 1). The capsid shell spans residues 157-336 which fold into a nine-helix bilobar CA domain structure, highly similar to Arc and Ty3 capsids (*18, 33*) (Supplemental Movie 2). Both the CA_NTD_ and CA_CTD_ of mPNMA2 are compact domains with a hydrophobic core and hydrophilic surface. The 28 residues C-terminal to CACTD are located on the inside of the capsid shell, whereas the 156 N-terminal residues protrude from the centre of the five-fold capsomers to form a disordered spike (Figure 1e; Supplemental Figure S7). The protein-protein packing within the mPNMA2 capsid is highly similar to the Ty3 retrotransposon capsid and the mature HIV-1 capsid, confirming the conservation of Gag capsid-assembly (Supplemental Figure S8). However, unlike dArc1, mPNMA2 does not contain any conserved nucleic acid binding motifs in the capsid interior part of the protein. The electrostatic potential of the interior capsid shell reveals only one exposed positively charged patch located at the twofold symmetry axis and this interface is likely transiently shielded by a highly negatively charged motif in the extended C-terminus of the protein (Supplemental Figure S9). These observations suggest that specific interactions with oligonucleotides are unlikely.

### PNMA2 is released from cells as non-enveloped capsids

Retrovirus capsids are released from cells as membrane-enveloped particles, similar to EVs. Arc protein is also released in EVs (*16*). To determine whether mPNMA2 is released from cells, we harvested media from primary cultured cortical neurons and fractionated the media using size-exclusion chromatography. mPNMA2 protein was isolated in early fractions that are also enriched with canonical EV proteins such as the ESCRT protein ALIX (Figure 2a). To determine whether released mPNMA2 protein is enveloped by a membrane, we performed a Proteinase K protection assay. Surprisingly, mPNMA2 protein was highly degraded by Proteinase K even without detergent present, unlike EV proteins, suggesting a lack of membrane protection (Figure 2b and c). These results contrasted with endogenous Arc protein release, which showed significant protection from Proteinase K (Figure 2b and c). To further determine whether mPNMA2 is released in EV fractions, we used iodixanol gradient ultracentrifugation and found that endogenous PNMA2 protein is enriched in a different fraction (fraction 6) to neuronal EV proteins (fraction 3), indicating released mPNMA2 differs in density from proteins found in canonical EV fractions (Figure 2d). Capsid structures were observed in iodixanol fraction 6, as determined by negative-staining EM (Figure 2e). To measure enough capsids to determine average size, myc-mPNMA2 was transfected HEK293T cells (Supplemental Figure S10). Capsids isolated from iodixanol fraction 6 and imaged on EM grids showed an average size similar to capsids from purified recombinant PNMA2 protein (Supplemental Figure S10e).

**Figure 2.**
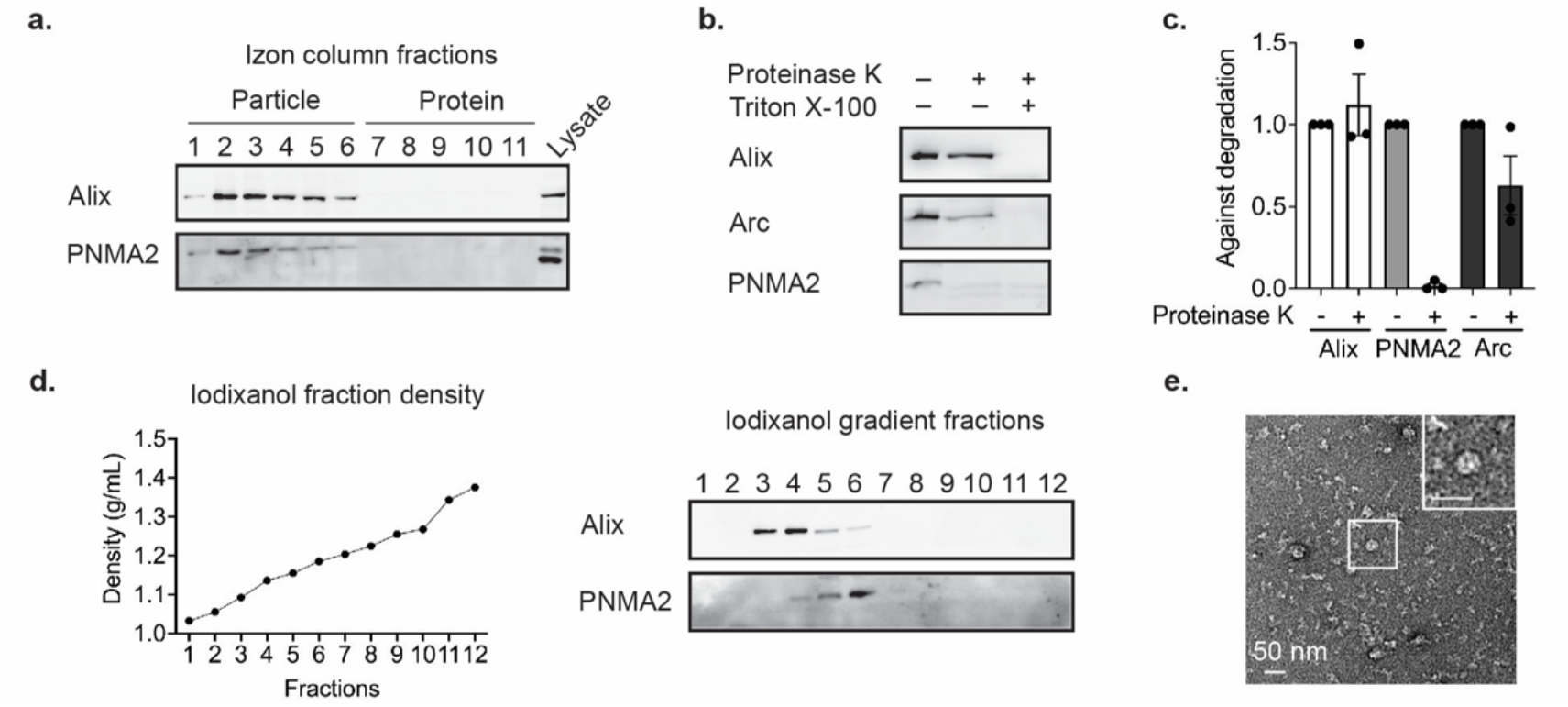
Endogenous mPNMA2 is released by neurons as non-enveloped capsids. **a**. Media was collected from primary cultured cortical neurons (DIV15-16) after 24h incubation and fractionated using size exclusion chromatography (SEC). Fractions were run on a gel and blotted for mPNMA2 protein and ALIX, a canonical EV marker. mPNMA2 protein is released in early fractions that contain EV proteins. **b**. The early fractions (1-4) from SEC were pooled and blotted for Arc, mPNMA2, and ALIX. Fractions were incubated with Proteinase K (200µg/mL) with or without detergent present (1% Triton-X) for 10mins. Representative Western blots show that mPNMA2 protein was sensitive to Proteinase K degradation without detergent present. **c**. Quantification of Western blot in b (n=3 experiments). Error bars indicate mean ± s.e.m. **d**. The early fractions (1-4) from SEC were fractionated using ultracentrifugation. An iodixanol gradient was used to separate proteins by density and size. mPNMA2 protein was enriched in fraction 6, while ALIX was enriched in fractions 3 and 4. **e**. Representative negative-stained EM image of non-enveloped mPNMA2 capsids isolated from iodixanol gradient fraction 6 in d.

Patients with PNMA2-related paraneoplastic syndrome often present with small cell lung cancer (*6*). To test whether tumor cells release hPNMA2, we used a human small cell lung cancer cell line (NCI-H378) that expresses a high level of hPNMA2 (*34*). We grew cells, harvested media after 24 hours, and performed size exclusion chromatography. hPNMA2 was released and present in early fractions, like ALIX (Supplemental Figure S11a). Similar to neurons, the released hPNMA2 was not protected from Proteinase K degradation (Supplemental Figure S11b) indicating release as non-enveloped capsids.

To determine whether mPNMA2 release requires assembled capsids, we designed point mutations predicted to disrupt the five-fold axis (Y162A – Y/A) or CA_NTD_-CA_CTD_ capsid interface (L270Q/L325Q – L/Q) between PNMA2 subunits (Supplemental Figure S7b and e) as mutations in the five-fold axis or CANTD-CACTD interface disrupt retrovirus capsid assembly (*35, 36*). As predicted, purified mPNMA2 Y/A and mPNMA2 L/Q proteins were unable to form capsids as analyzed by size exclusion chromatography, negative-stain EM, and mass photometry (Supplemental Figure S12 a-c). We overexpressed mPNMA2, mPNMA2 L/Q or mPNMA2 Y/A in HEK293T cells and measured protein release. The release of mPNMA2 L/Q and Y/A were significantly reduced compared with WT protein (Supplemental Figure S12 d-e). Together, these data show that mPNMA2 is released from cells as non-enveloped capsids.

### PNMA2 capsids induce autoantibody production

We next determined whether PNMA2’s virus-like properties are responsible for eliciting autoantibody production. Since PNMA2 appears to be exclusively expressed in neurons of the CNS (Figure 1b, Supplemental Figure 1), we hypothesize that aberrant expression of PNMA2 in tumor cells (*8-12, 37*) and release of capsids outside of the CNS may be immunogenic. Virus capsids induce a stronger immune response than soluble proteins due to the capsid’s influence on antigen transport, adaptive immune response, and cross-presentation (*38-41*). Thus, the ability of PNMA2 protein to form capsids that are released without membranes may explain its immunogenicity. To examine the immunogenicity of mPNMA2 capsids, we injected mice with 5µg purified recombinant mPNMA2 capsids without adjuvant and collected blood 3 weeks after injection. mPNMA2 capsid-injected mice produced high titers of antibodies that bind to mPNMA2 capsids, as assessed by ELISA (Figure 3a) and by immunogold EM (Figure 3b). The production of antibodies in mPNMA2 injected mice was also confirmed by Western blot (Supplemental Figure S13a). However, mice injected with 5µg of purified mPNMA2 L/Q capsid-mutant protein did not produce mPNMA2 L/Q-specific antibodies (Figure 3a, Supplemental Figure S13a). A second injection of 5µg mPNMA2 L/Q 3 weeks after the first injection, which may have primed the immune system, also did not induce autoantibody production further indicating a lack of immunogenicity (Supplemental Figure S13b). Capsid-induced antibodies showed similar endpoint titers against capsids and L/Q mutant protein when used as ELISA antigens, indicating that the lack of antibody signal in mPNMA2 L/Q injected mice is not due to a lack of exposed epitopes (Supplemental Figure S13c).

**Figure 3.**
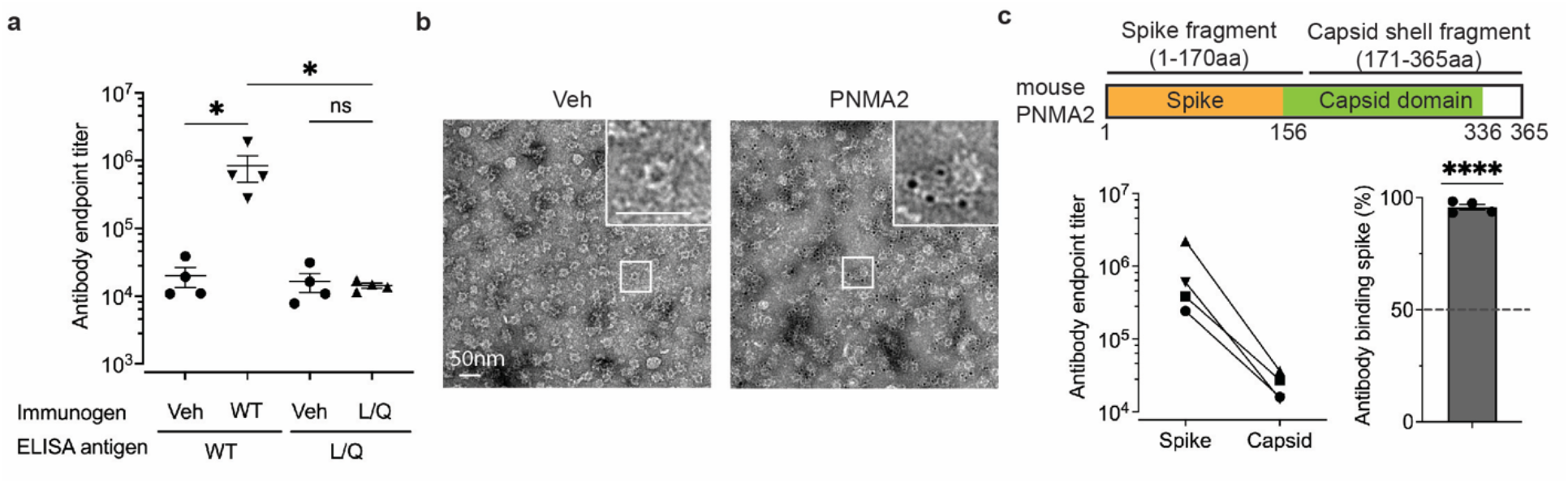
mPNMA2 capsids injected into mice induce PNMA2 autoantibody production. **a**. Mice were injected intraperitoneally with vehicle (n=4), 5µg purified mPNMA2 capsids (n=4) or 5µg mPNMA2 L/Q protein (n=4) and blood sera collected 3 weeks after injections. Sera were analyzed for antibody production using ELISA, using 2µg/mL mPNMA2 capsids or mPNMA2 L/Q protein as the antigen coated on the plates. mPNMA2 capsid-injected mice produced robust PNMA2 autoantibodies, whereas vehicle and mPNMA2 L/Q injections did not elicit autoantibody production. (*One-way ANOVA with post-hoc pairwise comparisons by Tukey’s test, P=0.0146. Vehicle vs. mPNMA2 WT (using mPNMA2 capsids as antigen): P=0.0308; Vehicle vs. mPNMA2 L/Q (using mPNMA2 L/Q as antigen): P>0.9999); mPNMA2 WT vs. mPNMA2 L/Q: P=0.0296.). **b**. Representative negative-stained EM images of purified mPNMA2 capsids immunogold labelled with mouse PNMA2 autoantibodies in serum collected from PNMA2 capsid-injected mice. **c**. Purified mPNMA2 spike fragments and capsid shell fragments (see schematic) were used as ELISA antigens to map the epitopes of PNMA2 autoantibodies from PNMA2 capsid-injected mice. The schematic shows PNMA2 protein regions located on the spike and the capsid domains of capsids, as determined by the cryo-EM structure. PNMA2 autoantibodies preferentially bind to the spike fragments. (****One sample t-test, P<0.0001, null hypothesis of 50% binding). Error bars indicate mean ± s.e.m. WT: wild type; L/Q: L270QL325Q; Veh: vehicle.

The structure of mPNMA2 capsids shows that the exposed “spikes” may be more immunogenic than the capsid body since these residues are more likely to be exposed to adaptive immune cells (*42, 43*). To test this, we purified the N-terminal region of mPNMA2 capsids (the “spike” aa 1-170) and the main capsid shell (aa 171-356) (Figure 3c). Autoantibodies from capsid-injected mice preferentially bind to the N-terminal “spike” fragment (Figure 3c).

To determine whether PNMA2 autoantibodies from human paraneoplastic patients bind hPNMA2 capsids, we obtained CSF samples from patients associated with paraneoplastic syndromes that have PNMA2 antibodies (see Table 1 for patient information) and control CSF samples (obtained from the Antineuronal Antibodies in Autoimmune Neurological Disease biorepository, University of Utah – IRB #00001919) that did not have PNMA2 autoantibodies (see Table 2). Using an ELISA coated with recombinant hPNMA2 capsids, we found that patient CSF contains high levels of PNMA2 autoantibodies capable of binding capsids (Figure 4a). PNMA2 specific antibodies in patient CSF also robustly labelled hPNMA2 capsids, as determined by immunogold EM (Figure 4b). Patient CSF PNMA2 autoantibodies preferentially bind to N-terminal “spike” epitopes predicted to be on the exterior surface of hPNMA2 capsids (Figure 4c), similar to autoantibodies produced in capsid-injected mice. These data strongly support the hypothesis that PNMA2 protein elicits an autoantibody response due to the high immunogenicity of PNMA2 virus-like capsids.

**Figure 4.**
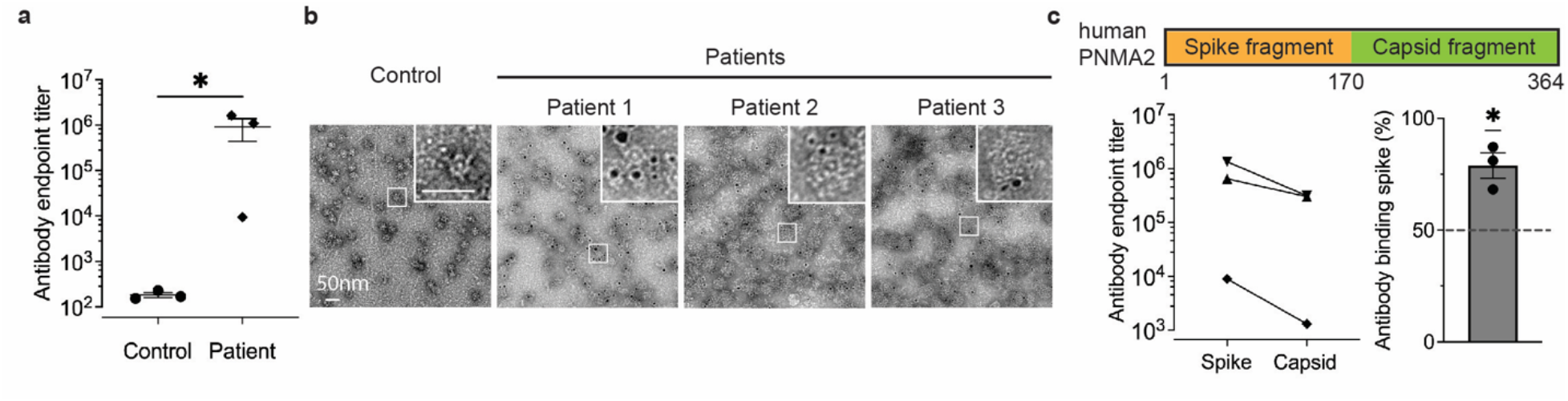
PNMA2 autoantibodies in CSF from human patients with paraneoplastic neurological syndrome preferentially bind to hPNMA2 capsids. **a**. An ELISA, using purified hPNMA2 capsids as the antigen, was used to quantify PNMA2 autoantibodies in CSF from patients diagnosed with or without PNMA2-related paraneoplastic neurological syndrome. (n=3 for each group, *Mann-Whitney test, P=0.05). **b**. Representative negative-stained EM images of hPNMA2 capsids labeled with immunogold using patient CSF from control or paraneoplastic patients. **c**. ELISAs using hPNMA2 spike and capsid shell fragments as antigens show patient CSF PNMA2 autoantibodies preferentially bind to the spike fragment (*One sample t-test, P=0.0356, null hypothesis of 50% binding). Error bars indicate mean ± s.e.m.

## Discussion

The discovery that Arc proteins form virus-like capsids intimated that other coopted Gag-derived proteins may also retain the ability to form capsids and that this property may have been repurposed repeatedly during evolution to function in intercellular communication (*14-16*). Other predicted Gag-like genes, including other *PNMA* genes, have also been shown to form virus-like particles (*14*). In this study we found that PNMA2 protein, which is highly expressed in mammalian cortex and hippocampus, forms small virus-like capsids. This indicates that there is a repertoire of mammalian endogenous virus-like capsids that have been evolutionarily re-purposed from retrotransposons (*26*).

In contrast to Arc release in EVs, we found that PNMA2 is released from cells as non-enveloped capsids. These capsids may be released via an unknown secretory pathway that could occur through non-canonical autophagy or lysosomal pathways (*44, 45*). Some viruses, such as adenoviruses and adeno-associated viruses (AAVs), are also released without membranes from cells. However, the cell biology and mechanisms of non-enveloped capsid release remain obscure. Since AAVs are used in current gene therapy applications and endogenous capsids may also be harnessed for gene delivery (*14*), understanding how non-enveloped capsids are released may facilitate gene therapy applications. However, our data caution that ectopic capsid expression may activate the immune system. The structural information we uncovered on PNMA2 capsids may provide a resource to design capsids that mitigate an autoimmune response.

Patients with PNMA2 autoantibodies experience neurological symptoms and signs, including cognitive impairment (*5*). Since normal expression of these genes is restricted to the brain, we posit that an antibody response may be elicited when tumors release high amounts of PNMA2 capsids outside the CNS. Whether autoantibody production correlates with overall PNMA2 levels or amount of release from tumors remains to be determined. The precise role of immune mechanisms in paraneoplastic neurological syndromes remain poorly characterized. However, prior brain pathologic analysis (autopsy or biopsy) demonstrating perivascular lymphocytic cuffing and interstitial infiltrates of lymphocytes with variable gliosis and neuronal degeneration (*7*). The inflammatory infiltrates were reported to be comprised mainly of T-cells, with smaller amounts of B-cells, macrophages and microglial activation, with a minority of samples also demonstrating plasma cells in the lymphocytic infiltrates (*7*). However, we find that PNMA2 autoantibodies in patient CSF can bind PNMA2 capsids, suggesting that these antibodies may also interfere with the putative intercellular signaling function of PNMA2. Capsids injected into mice induce a robust generation of autoantibodies that preferentially bind to external “spike” epitopes on capsids. However, mice injected with capsid disrupting mutant mPNMA2 protein showed no antibody response, highlighting the specific immunogenicity of capsids. Patient CSF PNMA2 autoantibodies also preferentially bind to external “spike” epitopes on PNMA2 capsids. We conclude that high levels of PNMA2 capsids found outside the CNS may induce a robust autoimmune response that may trigger a paraneoplastic neurological syndrome.

While PNMA2 is primarily and highly expressed in the mammalian brain, with conserved expression pattern in primates, the biological function of PNMA2 remains unknown. We have shown that PNMA2 protein forms virus-like capsids that are released from neurons and we speculate that these virus-like properties are important for PNMA2 function in the brain. Future studies will determine the function of PNMA2 capsids and whether other PNMA proteins form virus-like capsids that are released as non-enveloped particles. Our findings may help determine therapies for paraneoplastic neurological syndromes and more generally help reveal mechanisms of capsid-induced immune activation.

## Methods

### Plasmids

The coding sequence (CDS) of mPNMA2, hPNMA2, myc-mPNMA2 1-170aa, myc-mPNMA2 171-365aa, myc-hPNMA2 1-170aa, and myc-hPNMA2 171-364aa, mPNMA2 L270QL325Q were synthesized by Integrated DNA Technologies (Coralville, IA). These fragments were digested by *BamHI* and *XhoI* restriction enzymes (New England BioLabs, Ipswich, MA), and ligated into pGEX-6p1 vector for protein expression and purification. The mPNMA2 CDS was amplified by PCR, digested by *SalI* and *NotI*, and cloned into the PRK5-myc vector. Similarly, pLVX-myc-mPNMA2, pLVX-mPNMA2, pLVX-mPNMA2 L270Q/L325Q, and pLVX-mPNMA2 Y162A were constructed by PCR amplification of mPNMA2 WT and L270QL325Q CDS, *BamHI* and *XhoI* digestion, and ligation into pLVX backbone between *BamHI* and *XhoI* sites. All ligation products were transformed into NEB Stable Competent E. coli (High Efficiency) (New England Biolabs, Ipswich, MA). Individual colonies were inoculated to isolate plasmids and screen for correct constructs by sequencing.

### Protein purification

Bacterial protein expression plasmids were transformed into *E. coli* Rosetta2 strain (Novagen, Gibbstown, NJ) and the protein was expressed in ZY autoinduction media. *E. coli* were grown to 0.6-0.8 of OD600 at 37°C 150rpm, then switched to 19°C for 16-20h shaking. Afterwards, bacteria were pelleted down at 5000g for 15min at 4°C, resuspended in lysis buffer (500mM NaCl, 50mM Tris, 5% glycerol, 1mM DTT, complete Protease Inhibitor Cocktail, pH 8.0). The resuspended bacteria were frozen by liquid nitrogen and then thawed. The thawed lysates were sonicated and centrifuged at 21,000g for 45mins at 4°C. The supernatant was filtered through 0.45µm filters and incubated with Sepharose 4B beads (Cytiva, Marlborough, MA) overnight at 4°C. GST-tagged proteins were eluted from Sepharose beads by incubating with L-reduced glutathione (20mM, pH 8.0) and exchanged to cleavage buffer (50mM Tris, 150mM NaCl, 1mM DTT, 1mM EDTA, pH 7.2) using a Vivaspin column (Cytiva, Marlborough, MA). The cleavage of GST was performed by PreScission Protease (Cytiva, Marlborough, MA) overnight at 4°C. The cleaved PNMA2 protein was run through a HiLoad 16/600 Superdex 200 pg (Cytiva, Marlborough, MA) size-exclusion column in the capsid assembly buffer (500mM phosphate, 50mM Tris, 0.5mM EDTA, pH 7.5) to further purify PNMA2 and facilitate capsid formation. For protein purified for mice immunization, Triton X-114 was used to wash GST-tagged protein bound to Sepharose beads to remove endotoxin.

### Negative-staining electron microscopy

Copper 200-mesh grids coated with formvar and carbon (Electron Microscopy Sciences, Hatfield, PA) were discharged for 30s in a vacuum chamber. Grids were incubated with 3.5µL purified protein (0.5mg/mL) for 45s and excess sample was wicked away. Grids were then immediately washed twice with 30uL water for 5 s and once with 15µL 1% uranyl acetate (UA) for 5s. Excess UA was wicked away and the grids were stained with 15µL 1% UA for 30s. Excess UA was wicked away and grids were air-dried. The grids were imaged using a FEI Tecnai T12 Transmission Electron Microscope operated at 120kV.

### Cryo-electron microscopy

4 μl of mPNMA2 capsids at a concentration of ∼1 mg/ml were applied to glow-discharged QUANTIFOIL 2/2 Au continuous carbon grids. The sample was blotted and plunge-frozen in liquid ethane using a Vitrobot Mark IV. Cryo-EM images were acquired on a 300 keV FEI Titan Krios microscope with X-FEG emitter equipped with a Falcon III detector operated in counting mode. mPNMA2 datasets were collected using EPU (ThermoFisher) at a nominal magnification of 96000x with a calibrated pixel sizes of 0.832 Å. Micrographs were recorded as movies divided into 75 frames with a final accumulated dose of 40 e/Å^2^. Defocus values ranged from −0.5 to −2.5 μm.

### Image processing

mPMNA2 image processing and reconstructions are summarized in Supplemental Figure S5 and S6 and Supplemental Table S1. Acquired movies were aligned using 5×5 patches, averaged and dose-weighted using RELION4 (*46*). Contrast transfer function (CTF) parameters were estimated using CTFFIND4 (*47*). Particles were automatically picked using a retrained topaz model (*48*). Boxsize used for extraction was 512×512 pixels. Extracted particles were subjected to several rounds of 2D classification to remove false picks and junk particles. 2D classes showed views characteristic for icosahedral symmetry, and icosahedral symmetry (I) was applied for initial model generation and throughout subsequent 3D classification and refinement. Finally, we performed per-particle CTF estimation, Bayesian polishing and Ewald sphere correction. The effective resolutions of the cryo-EM density maps were estimated by Fourier shell correlation (FSC = 0.143) according to the definition of Rosenthal and Henderson.

To further improve the resolution, we performed symmetry expansion as implemented in RELION4, to calculate the positions and orientations for each of the asymmetric units centered either at the five, three or two-fold symmetric axes. We extracted individual capsomeres using box sizes of 220×220 pixels. We performed 3D classification without alignment to remove bad views and to select classes for subsequent refinement. For all maps, local resolutions were calculated using ResMap (*49*) and maps were locally sharpened using Phenix (*50*).

### Model building

For model building we used Alphafold2 (*51*) to generate an initial model. After initial rough fitting using chimera (*52*), we went through several rounds of manual and auto-refinement in coot, Phenix, and ISOLDE (*53, 54*). Model statistics are shown in Supplemental Table S1.

### Mass photometry

Mass photometry was performed on a Refeyn TwoMP mass photometer. Protein stocks were brought to room temperature and diluted with stock buffer (500mM Na2HPO4, 50mM Tris, 10% glycerol, pH 7.5) to a concentration of 100nM. A 20□l droplet of each sample dilution was placed on top of a clean chambered-coverslip mounted on the oil objective of the instrument. Following autofocus stabilization, a one-minute movie was recorded using the Aquire-MP (Refeyn) software. A protein standard containing equimolar amounts of four proteins (conalbumin, aldolase, ferritin and thyroglobulin) with a molecular mass ranging from 75 to 669 kDa was also measured and used for calibration. Movies were converted to mass using a contrast-to-mass calibration and analyzed using the Discover-MP software (Refeyn).

### Primary cortical/hippocampal neuronal culture

Primary neuron cultures were prepared from male and female E18 WT mouse cortex and hippocampus as previously described (*55*). Tissue was dissociated in DNase (0.01%; Sigma-Aldrich, St. Louis, MO) and papain (0.067%; Worthington Biochemicals, Lakewood, NJ), and then triturated with a fire-polished glass pipette to obtain a single-cell suspension. Cells were pelleted at 500xg for 4 min, the supernatant removed, and cells resuspended and counted with a TC-20 cell counter (Bio-Rad, Hercules, CA). Neurons were plated on glass coverslips (Carolina Biological Supply, Burlington, NC) coated with poly-l-lysine (0.2 mg/mL: Sigma-Aldrich, St. Louis, MO) in 12-well plates (Greiner Bio-One, Monroe, NC) at 90,000 cells/mL, or in 10-cm plastic dishes at 800,000 cells/mL. Neurons were initially plated in Neurobasal media containing 5% serum, 2% GlutaMAX, 2% B-27, and 1% penicillin/streptomycin (Thermo Fisher Scientific, Waltham, MA) in a 37°C incubator with 5% CO2. On DIV4, neurons were fed via half media exchange with astrocyte-conditioned BrainPhys™ Neuronal Medium (Stemcell Technologies, Vancouver, Canada) containing 1% serum (Thermo Fisher Scientific, Waltham, MA), 0.25% L-Glutamine (Thermo Fisher Scientific, Waltham, MA), 1% penicillin/streptomycin (Thermo Fisher Scientific, Waltham, MA), 2% SM1 (Stemcell Technologies, Vancouver, Canada), and 5 μM cytosine β-d-arabinofuranoside (AraC) (Sigma-Aldrich, St. Louis, MO). Half media exchange of astrocyte-conditioned media was conducted every three days thereafter.

### HEK 293T cell culture

HEK 293T cells (#CRL-11268) were purchased from ATCC. Cells were cultured at 37°C with 5% CO2 in media including DMEM (Thermo Fisher Scientific, Waltham, MA), 5% fetal bovine serum (Thermo Fisher Scientific, Waltham, MA) and 1% penicillin/streptomycin (Thermo Fisher Scientific, Waltham, MA). Cell cultures were passaged at 70% confluency.

### NCI-H378 cell culture

NCI-H378 cells (#CRL-5808) were purchased from ATCC. Cells were cultured at 37°C with 5% CO2 in media including RPMI-1640 (ATCC, Manassas, VA), 5% fetal bovine serum (ATCC, Manassas, VA), and 1% penicillin/streptomycin (Thermo Fisher Scientific, Waltham, MA).

### HEK 293T cells transfection

10 cm dish transfection: 9µg of plasmids and 25µL 1µg/µL Polyethylenimine (Polysciences, Warrington, PA) were mixed in 1mL Opti-MEM (Thermo Fisher Scientific, Waltham, MA) and incubated at room temperature for 20 minutes. This mixture was added to 60% confluent cell culture in transfection media (DMEM+10% FBS) and incubated overnight.

### LDH cytotoxicity assay

The amount of LDH in the cell lysates and media was measured using CyQUANT LDH Cytotoxicity Assay kit (Thermo Fisher Scientific, Waltham, MA). The percentage of cell death was calculated by the total amount of LDH in the media divided by that in the cell lysates.

### Lentivirus production and transduction

HEK293 cells in a 10 cm dish were transfected with pLVX-VSVG, pLVX-psPAX 2, and pLVX-GFP or pLVX-myc-mPNMA2 overnight. Transfection media was changed to regular culture media (DMEM+10% FBS+ 1% penicillin/streptomycin). The media was collected 24 hours later and centrifuged at 1,000g for 10min. 20mM HEPES was added to the supernatant and centrifuged at 5,000g overnight. Lentivirus particles were pelleted and resuspended in 400 µL PBS. Cultured neurons in a 10 cm dish were transduced with 200 µL lentivirus particles at DIV13 for 2 days.

### Mice immunization

10–12 week-old mice (C57BL/6) were used for immunization. Mice were injected intraperitoneally with vehicle (capsid assembly buffer), 5µg of mPNMA2 capsids, or 5µg of mPNMA2 L270QL325Q capsid mutant on day 0. The immunized proteins were not mixed with any adjuvant. Blood was collected on day 21 via the tail vein. A second injection of 5µg of mPNMA2 L270QL325Q capsid mutant and vehicle were performed on day 21 and blood was collected on day 42. Blood was coagulated at 4°C overnight and serum was harvested by centrifuging twice at 2,000g for 10min to collect the supernatant. Endotoxin was removed from the purified protein by Triton X-114 wash. Preparations were checked for the presence of capsids via negative stain EM, prior to injection.

### Extracellular vesicle and capsid purification from culture media

For HEK 293T cells, full media change was performed 12 hours after transfection and then collected 24 hours later. For neuronal culture, full media change was performed at DIV15 (2 days after lentivirus transduction) and then collected 24 hours later. Collected media was centrifuged at 2,000g for 10min and then 20,000g for 20min. The supernatant of the collected media was concentrated into 0.5mL using Vivaspin columns. Concentrated media was loaded onto IZON mini-size exclusion columns (IZON Sciences, Christchurch, New Zealand). The first 3mL run through the column was collected as the void fraction. Then, every 0.5mL was collected for a total of 11 fractions, labeled as fraction 1 to 11.

### Proteinase K protection assay

Cultured media was processed using IZON columns as above and treated under three different conditions: 1. 20µL sample + 7µL H2O. 2. 20µL sample + 6µL H2O + 1µL proteinase K (200µg/mL) (New England BioLabs, Ipswich, MA). 3. 20µL particle sample + 3µL H2O + 3µL 10% Triton X-100 + 1µL proteinase K (200µg/mL). Media was incubated at room temperature for 10 min. Then, 3µL of 10mM PMSF was added and incubated for 10min at room temperature. Finally, samples were mixed with laemmli buffer (40% glycerol, 250 mM Tris, 4% SDS, 50 mM DTT, pH 6.8) and boiled at 95°C for 5 min.

### Western blot

Samples were mixed with 4x laemlli buffer and incubated at 95°C for 5 mins. Proteins were loaded and separated by 10% or 12% SDS-PAGE gel, followed by wet transfer to a nitrocellulose membrane (GE Healthcare, Chicago, IL). Total protein was stained and destained by Pierce Reversible Protein Stain Kit (Thermo Fisher Scientific, Waltham, MA). The membrane was blocked by 5% milk in TBS for 1h at room temperature. Primary antibodies were diluted in 1% milk in TBS and incubated with the membrane at 4°C overnight. Primary antibodies include: anti-myc (05-724, Sigma-Aldrich, St. Louis, MO; 1:1000), anti-PNMA2 (16445-1-AP, Proteintech, Rosemont, Illinois; 1:1000), anti-Alix (customized antibody from Wesley Sundquist’s lab; 1:500), mice sera (1:1000). The membrane was washed by TBS and then incubated with secondary antibodies (anti-mouse IgG, anti-rat IgG, anti-human IgG, Jackson Laboratory, Bar Harbor, ME; 1:5000) diluted by 1% milk in TBS for 1h at room temperature. Afterwards, the membrane was washed with TBS for three times. Bound antibodies were detected by Clarity™ Western ECL Substrate (Bio-Rad, Hercules, CA) and imaged using an Amersham ImageQuant™ 800 Western blot imaging systems (Cytiva, Marlborough, MA). Images were analyzed and quantified using ImageJ.

### RNA-Scope

Brian tissues obtained from 2-month-old mice and primary hippocampal neuronal cultures (DIV15) were used for RNAscope. RNAscope multiplex fluorescent v2 assay kit and the probe were purchased from Advanced Cell Diagnostics (Hayward, CA). The probe was tagged to opal 570 dye from Akoya Biosciences (Marlborough, MA).

### Confocal microscope imaging

Coverslips were imaged using a 10x objective for brain tissues and 60x oil objective for cultured neurons on a Nikon FV1000 confocal microscope (Tokyo, Japan). The images were analyzed using ImageJ software (National Institutes of Health, Bethesda, MD).

### Iodixanol gradient ultracentrifugation

An Iodixanol gradient was made with 2mL 15% (1M NaCl), 3mL 30%, 3mL 40% and 3mL 50% OptiPrep™ iodixanol (Alere Technologies AS, Oslo, Norway). 500µL of the sample was added to the top of the gradient and centrifuged at 280,000 rpm for 48 hours. After ultracentrifugation, sequential 1mL collections were made from the top of the gradient and labeled as fractions 1 to 12.

### Immunogold EM

Copper 200-mesh grids coated with formvar and carbon (Electron Microscopy Sciences, Hatfield, PA) were discharged for 30s in a vacuum chamber. PNMA2 purified protein (0.1mg/mL, 3.5µL) was loaded onto the grid and incubated for 45s. Grids were washed twice with 30uL diH2O for 30s and blocked with 30uL 1% BSA for 30mins. Grids were then incubated with sera/CSF/primary antibody in 30uL 1% BSA for 1h at room temperature and then washed three times with 30uL 1% BSA for 30s. Grids were then incubated with 6 nm immunogold conjugated secondary antibody, including Goat-anti-Human IgG (H&L) and Goat-anti-Mouse IgG (H&L) (Electron Microscopy Sciences, Hatfield, PA) (1:20 dilution in 1% BSA, 30uL) for 1h at room temperature. Afterwards, grids were washed three times with 30uL PBS for 30s and fixed with 30uL 2% glutaraldehyde for 5mins. Finally, grids were washed with 30uL diH20 for three times and stained with 30uL 1% uranyl acid for 1 min. The grids were imaged using a FEI Tecnai T12 Transmission Electron Microscope operated at 120kV.

### Enzyme-Linked Immunosorbent Assay (ELISA)

ELISAs were used to determine the titer and binding affinity of antibodies in human CSF and mice sera to different antigens. Maxisorp plates (Thermo Fisher Scientific, Waltham, MA) were coated overnight with 2µg/mL of purified PNMA2 capsids or PNMA 2 L270QL325Q protein in assembly buffer at room temperature. PNMA2 fragments were coated in PBS at 4°C. After coating, plates were washed with wash buffer (Thermo Fisher Scientific, Waltham, MA) 5 times and then blocked with 5% Bovine Serum Albumin (BSA; Sigma-Aldrich, St. Louis, MO) in PBS for 2 hours at 37°C. CSF and sera were diluted serially in 1% BSA and incubated with the plates for 2 hours at 37°C. After washing, plates were incubated with anti-mouse IgG or anti-human IgG (1:5000 in 1% BSA; Jackson Laboratory, Bar Harbor, ME) for 1 hour at 37°C. Afterwards, plates were washed and incubated with 3,3’,5,5’-Tetramethylbenzidine (TMB) substrate (Thermo Fisher Scientific, Waltham, MA) for 30 mins at 37°C. The enzyme reaction was stopped by adding the stop buffer (Thermo Fisher Scientific, Waltham, MA) and the absorbance was measured at 450nm.

### Quantification and statistical analysis

Data were analyzed using GraphPad Prism. Mann-Whitney test, paired t-test, or two-tailed t-test were used for the analysis. Not significant (ns) p>0.05; *p<0.05; **p<0.01; ***p<0.001; ****p<0.0001.

### RNA-seq data analysis

We obtained the processed single-cell expression matrix (counts) from Allen’s brain map (https://portal.brain-map.org) representing the human cortex and mouse cortex-hippocampus. We then used Seurat (v3.1.1) (https://github.com/satija.lab/seurat) within the R environment (v3.6.0) for processing the dataset. We kept the cells with minimum and maximum of 1,000 and 5,000 genes expressed (≥1 count), respectively. Moreover, cells with more than 5% of counts on mitochondrial genes were filtered out. The data normalization was achieved by scaling it with a factor of 10,000, followed by natural-log transformation. Cell type assignment was performed based on the annotations provided by the original publication, albeit we grouped the clusters into broader lineages of excitatory, inhibitory, Oligodendrocyte precursor cells (OPC), Oligodendrocytes and Glial cells (Astrocytes and Microglia). All the given annotations were further confirmed by their respective markers. The expression levels of each gene in a cluster correspond to the average log2 expression level scaled to the number of unique molecular identification (UMI) values captured in single cells. Finally, the sample replicates for each gene were aggregated per cell type, and their expression was calculated as trimmed means. For the analysis of Human, Chimpanzees and Macaque, we downloaded the human brain transcriptome from 33 major regions representing four adult healthy human individuals, three chimpanzees, and three rhesus macaques (GSE 127898). Trimmed mean of M (TMM) values normalized transcripts per million (TPM) values were imported in R, and variable features were identified when the ratio of variance and mean expression values were greater than one.

### Code availability

All statistical and plotting details, including the statistical tests, used, and precision measures for RNA-seq data analysis can be found on the GitHub repository at https://github.com/Manu-1512/MamGyp-int-rides-PNMA2

## Acknowledgments

We thank Wesley Sundquist and John McCullough for helpful discussions and experimental support. We thank Tammy Smith for helping provide human CSF samples. We thank Jenifer Einstein and Michael Hantak for generating primary cultured neurons, and all members of the Shepherd lab for technical help and support.

## Author contributions

J.X performed the RNAscope, biochemistry, electron microscopy, and ELISA experiments. S.E conducted the cryo-EM experiments. M.S performed phylogenomic and RNA-seq analysis. M.R made the slices for RNAscope and conducted the mice immunization and blood collection. I.I performed mass photometry experiments. G.S.D, A.L.P, and S.L.C provided the clinical human samples. J.X, S.E, M.S, M.R, C.F, J.A.G.B, and J.D.S. conceived and designed experiments. J.X and J.D.S. wrote the manuscript; all authors discussed results and edited the manuscript.

---

**Table 1 - Patient sample information**

## Patient Sample 1, *PNMA2 positive in CSF*

27-year-old previously healthy male presented with a 2-month history of lethargy and decreased libido. Neurological examination demonstrated normal cognition, with mild dysarthria, bilateral ptosis, slowing of upward saccades, and decreased arm swing. Screening blood work confirmed low testosterone, follicular stimulating hormone, and luteinizing hormone. Brain MRI disclosed an enhancing mass in the tectal region with dilation of the ventricles. Cerebrospinal fluid analyses (CSF) showed pleocytosis (13 cells per mm3; reference <6 cells per mm3) with lymphocytic predominance, elevated protein (58 mg/dL; reference 15-45 mg/dL), and normal glucose. There were no unique oligoclonal bands. CSF infectious studies were negative including herpes simplex virus polymerase chain reaction. CSF cytology and cytometry were negative for malignant cells. No disease-associated autoantibodies were detected on serum or CSF screening (Mayo Clinical Laboratories, “autoimmune encephalitis panel”, and anti-aquaporin4 antibodies). Stereotactic-guided tectal biopsy was performed, with pathology confirming a lymphoplasmacytic inflammatory process. Over the ensuing two months, the patient complained of increasing fatigue / lethargy, with double vision attributed to vertical gaze restriction. Repeat neuroimaging showed mild interval improvement in the tectal mass, with normal sized ventricles, with emergence of bilateral T2-FLAIR signal within the temporal lobes. Atypical paraneoplastic causes were considered, and body imaging was pursued. Body CT demonstrated numerous subcentimeter nonspecific nodes in the abdomen and pelvis. Testicular ultrasound confirmed bilateral testicular microlithiasis. A diagnosis of ma1/ma2 autoimmune encephalitis was considered and confirmed through testing of CSF and serum via Athena Diagnostics. Bilateral orchiectomy was performed given concern of disease-associated tumor. Histopathology was notable for dystrophic calcifications and fibrosis, suggestive of a burnt-out germ cell tumor. High-dose intravenous steroids were provided, with a corresponding brief improvement in alertness and temporal lobe T2-hyperintensities. Long-term follow-up was complicated by treatment-refractory temporal lobe seizures, necessitating escalation of immunotherapy, including monthly intravenous cyclophosphamide. Severe fatigue, lethargy, double vision, and vertical gaze palsy persisted.

## Patient Sample 2, *PNMA1 (Ma1) and PNMA2 (Ma2) positive in CSF*

54-year-old female with history of renal cell carcinoma status post-resection presented with a 6-month history of progressive diplopia, gaze paresis, abnormal gait with falls, increasing confusion, and new-onset seizures. On exam, the right eye was deviated downward, adduction was impaired in the left eye, and gait was wide-based and apractic. Brain MRI was markedly abnormal, with symmetric subcortical T2-hyperintensities involving the medial temporal lobes, parietooccipital cortices and medial thalami, concerning for posterior reversible encephalopathy syndrome versus neuroinflammation. CSF analyses disclosed 5 white blood cells (per mm^3^), normal glucose, and protein, with 9 CSF-specific bands. Noting the past history of renal cancer, body imaging was performed leading to detection of mediastinal and bilateral hilar lymphadenopathy, with bilateral pulmonary nodules and masses and a left adrenal mass. Lymph nodes were FDG-avid on follow-up PET scan. Mediastinal biopsy confirmed metastatic renal carcinoma. Paraneoplastic panel (Mayo Clinic Laboratories) including Ma1/Ma2 testing (Athena Diagnostics) identified Ma1/Ma2 antibodies in blood and CSF. High-dose IV methylprednisone and plasmapheresis (5 exchanges) were provided for treatment of Ma1/Ma2 paraneoplastic encephalitis without improvement in symptoms. The patient developed a diencephalic syndrome with ophthalmoparesis (near plegia) with “sunsetting eyes”, progressive encephalopathy, and cortical blindness. Planned treatment of her underlying malignancy was limited by the development of bilateral pulmonary emboli requiring systemic anticoagulation, and a general decline in physical health. The patient died of complications of her illness.

## Patient Sample 3, *PNMA1 (Ma1) and PNMA2 (Ma2) positive in CSF at > 1:32*

66-year-old male with a history of chronic tobacco use presented with two months of worsening balance, diplopia, and confusion. He had no prior medical conditions. Neurological exam revealed short-term memory deficits, a poor fund of knowledge, ataxia, opsoclonus, restricted upgaze, restriction of right eye abduction, and a right extensor toe response. Brain MRI revealed an encephalitis involving the brainstem, mesial temporal lobes, and basal ganglia. Cerebrospinal fluid (CSF) analysis showed a pleocytosis of 75 nucleated cells per mm^3^ (reference < 6 nucleated cells per mm^3^) with a lymphocytic predominance, elevated protein of 75 mg/dL (reference 15–45 mg/dL), and normal glucose. Five unique oligoclonal bands were present. *PNMA1 (Ma1) and PNMA (Ma2)* were positive (titer > 1:32, Athena Diagnostics). Low positive Anti-NMDA-R antibody was detectable only on a neat, undiluted cell-based assay (Mayo Clinical Laboratories), and serum studies demonstrated low positive anti-GAD65 antibodies at a titer present in up to 8% of the normal population and not generally consistent with symptoms or associated neurologic disease (0.12 nmol/L, reference ≤0.02), with the rest of the serum autoimmune encephalopathy evaluation otherwise negative (Mayo Clinical Laboratories). CSF infectious studies were negative including herpes simplex virus polymerase chain reaction. CSF cytology and cytometry were negative for malignant cells. Malignancy screening included a negative testicular ultrasound and a computed tomography (CT) of the chest that demonstrated extensive pulmonary fibrosis, several prominent mediastinal lymph nodes, and multiple segmental pulmonary emboli. A body positron emission tomography CT demonstrated FDG uptake in a right hilar lymph node in addition to a subpleural, sub-centimeter pulmonary nodule in the left upper lobe, but a progressive deterioration of his respiratory status prevented full investigation.

**Table 2 - Control CSF patient information**

## Control sample 1

60-year-old male with 2 years of progressive spasticity (bilateral lower extremities, progressed to left upper extremity). Neurologic exam notable initially for spasticity and upper motor neuron signs, but no lower motor neuron signs. Initial EMG with no abnormal spontaneous activity and MRI brain, C-, T-spine unremarkable. CK mildly elevated at 304. CSF with 2 WBC (ref 0-5), prot 52 (ref 14-45), glc 61 (ref 50-80), OCB neg, <0.0 IgG synthesis rate. Mayo autoimmune encephalopathy evaluation of serum and CSF with no informative autoantibodies detected. Glycine receptor antibody also negative in serum (not tested in CSF). Wash University neuromuscular antibody panel on serum also unremarkable. Ultimately progressed, and EMG findings with upper and lower motor neuron findings; diagnosed with amyotrophic lateral sclerosis.

## Control sample 2

54-year-old male followed for postherpetic neuralgia. CAF with WBC 0, prot 38, glc 65, OCB 0, <0.0 IgG synthesis rate. CSF culture negative, HSV1/2 Ab screen IgG <0.34, HSV Type 1 Ab IgG 0.0, HSV 1 and or 2 Abs IgM 0.19, VZV IgG <10, VZV IgM 0.02, VZV PCR not detected, CMV IgM <8, HSV 6 Ab IgM by IFA <1:20. Note-all these results are within normal limits. No send out antibody testing done because no concern for immune-mediated process.

## Control sample 3

58-year-old female with diffuse paresthesia and pain, diagnosed with small fiber neuropathy, pernicious anemia and positive GAD65 antibody (without neurologic manifestations). CSF WBC 5, prot 37, glc 53, OCB negative (note: matched bands in serum and CSF), IgG synthesis rate <0.0, Mayo Paraneoplastic Autoantibody Evaluation from CSF positive for GAD65 Ab assay 0.04 nmol/L (ref 0.02), negative for AGNA1, Amphiphysin Ab, ANNA1, ANNA2, ANNA3, CRMP5 IgG, PCA1, PCA2, PCA-Tr. Mayo Autoimmune dysautonomia eval (serum) with AChR Ganglionic Neuronal Ab 0.03 nmol/L (ref <0.02), GAD65 Ab 20.1 nmol/L (ref <0.02), negative for ANNA1, Striational Ab, N-type Calcium Channel, Ach Receptor binding Ab, Neuronal VGKC Ab, P/Q Type Calcium channel. Intrinsic factor blocking antibody positive. GAD65 Ab at ARUP >250 IU/mL (ref 0-5)

**Supplemental Figure S1:**
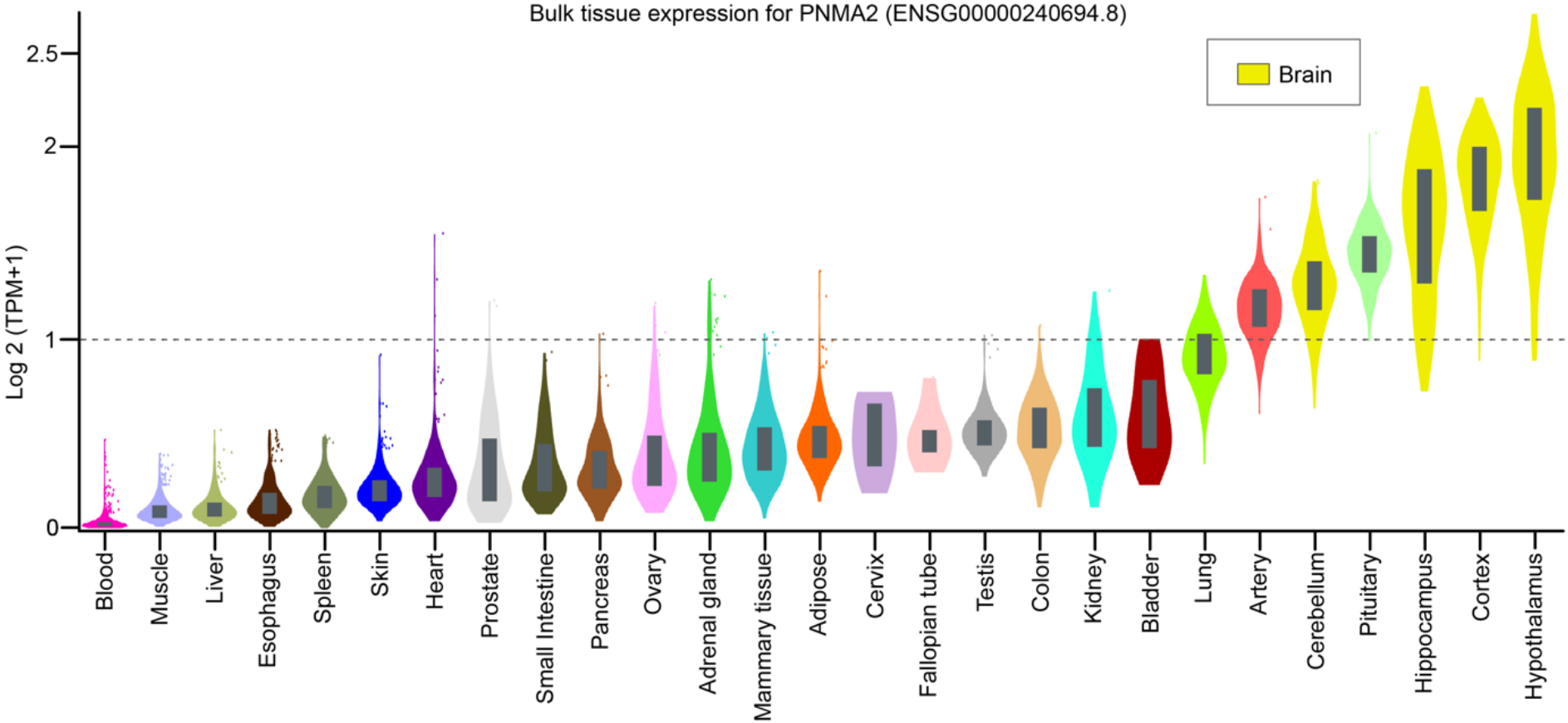
RNA expression of PNMA2 in bulk tissues of humans. Violin plots show the level of bulk RNA expression (Log2 (TPM+1)) of *PNMA2* gene from various adult tissues covered and analyzed by the Gtex consortium (https://www.gtexportal.org/home/gene/ENSG00000240694). Multiple tissues from the same organ were aggregated in one to simplify the visualization.

**Supplemental Figure S2.**
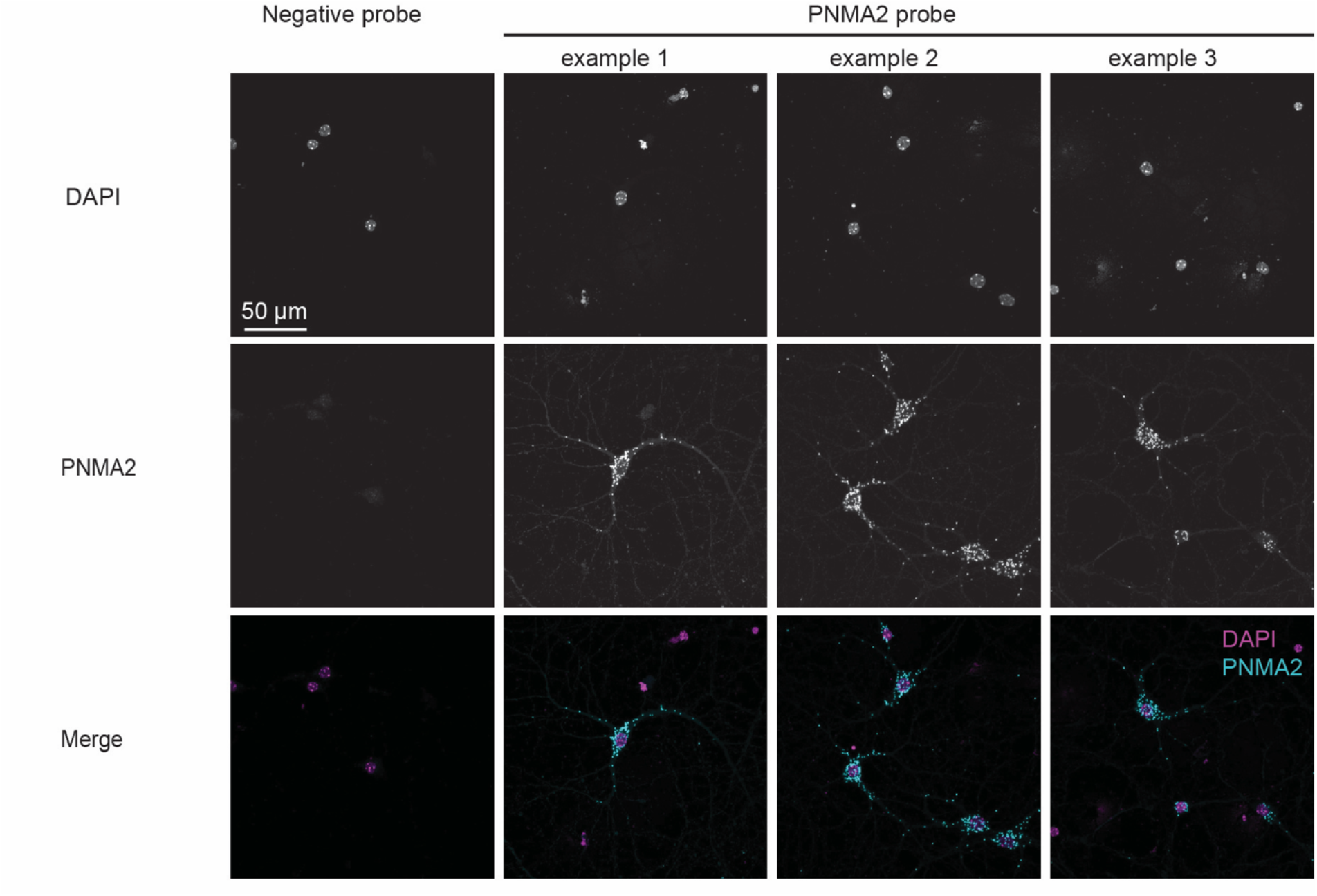
PNMA2 is highly expressed in hippocampal neurons. PNMA2 probe and negative probe were used to detect PNMA2 RNAs in the wild-type mouse hippocampal neuronal culture (DIV15) by RNAscope.

**Supplemental Figure S3:**
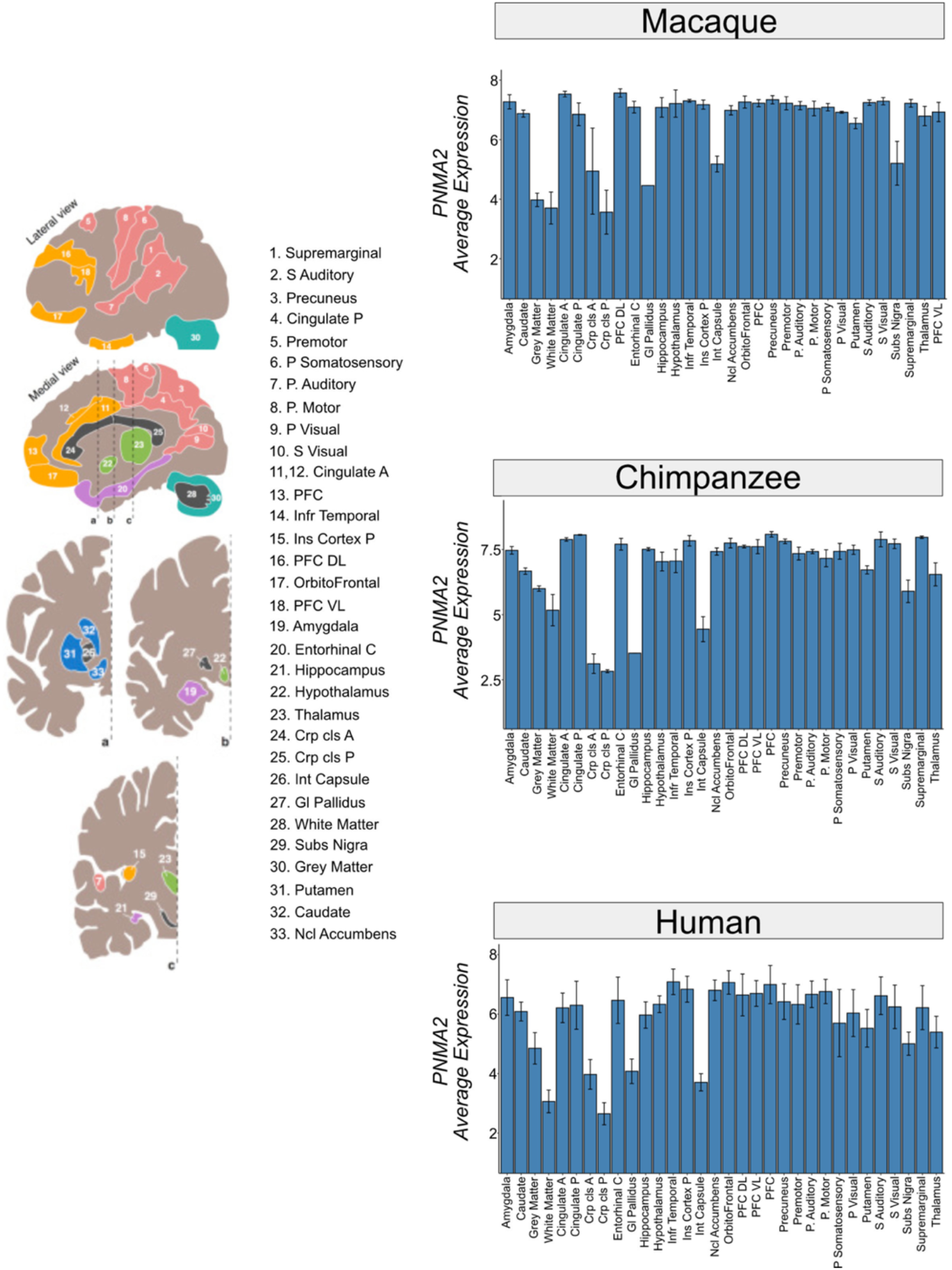
RNA expression of *PNMA2* and *DPYSL2* in brain regions of different primates. Combined bar plots show the level of RNA expression of *PNMA2* and *DPYSL2* genes at the log2 transcript per million (TPM) scale from the major structures of 33 brain regions by the analysis of 422 polyA-selected RNA-seq datasets in total. The data represents four adult healthy humans, three chimpanzees, and three rhesus macaques. Error bars show the mean +/- standard error of the replicates between individuals.

**Supplemental Figure S4.**
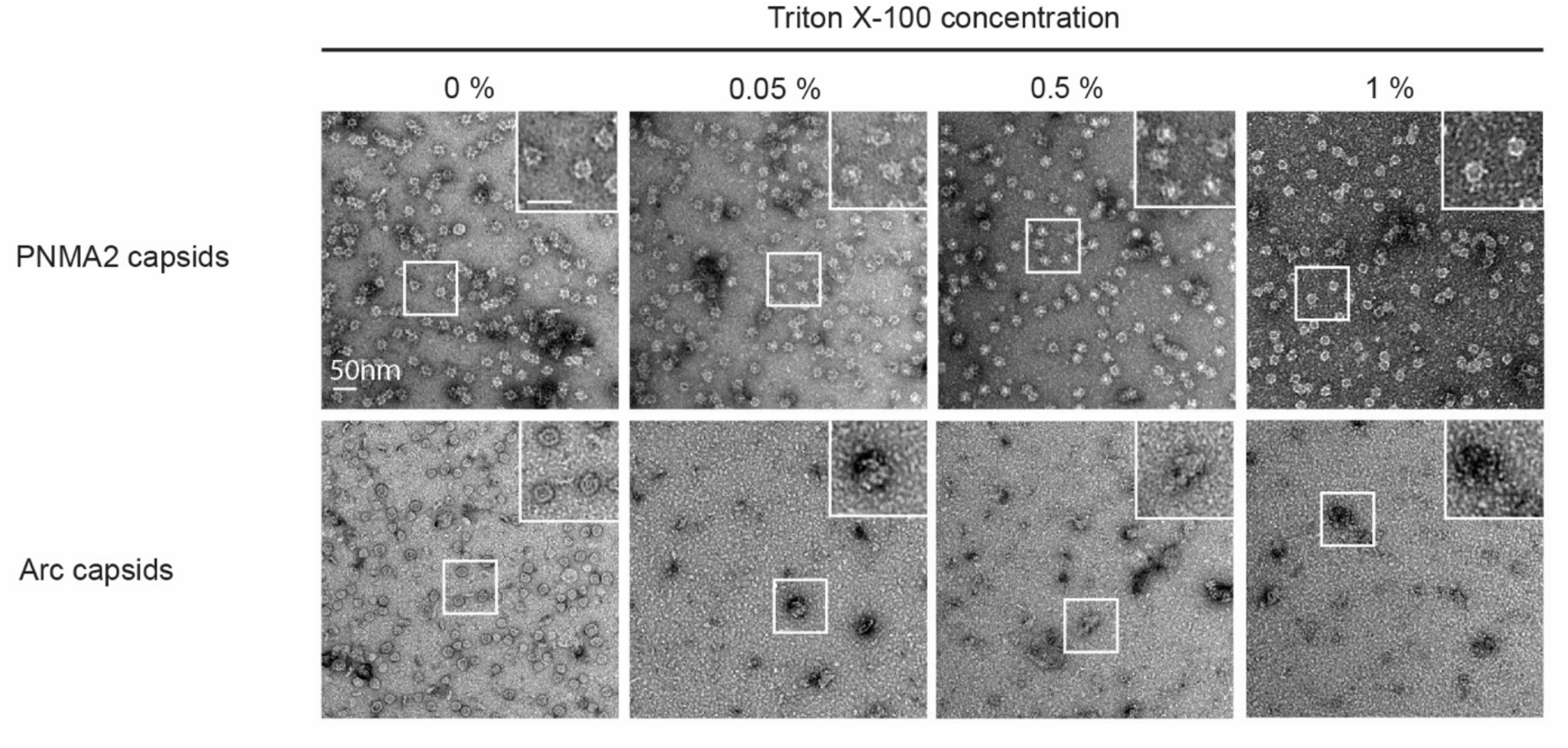
mPNMA2 forms stable capsids that are resistant to detergent. Representative negative-staining EM images of purified mPNMA2 and Arc capsids incubated with different concentrations (0%, 0.05%, 0.5%, 1%) of Triton X-100. Negative control: neuronal protein Arc formed capsids.

**Supplemental Figure S5:**
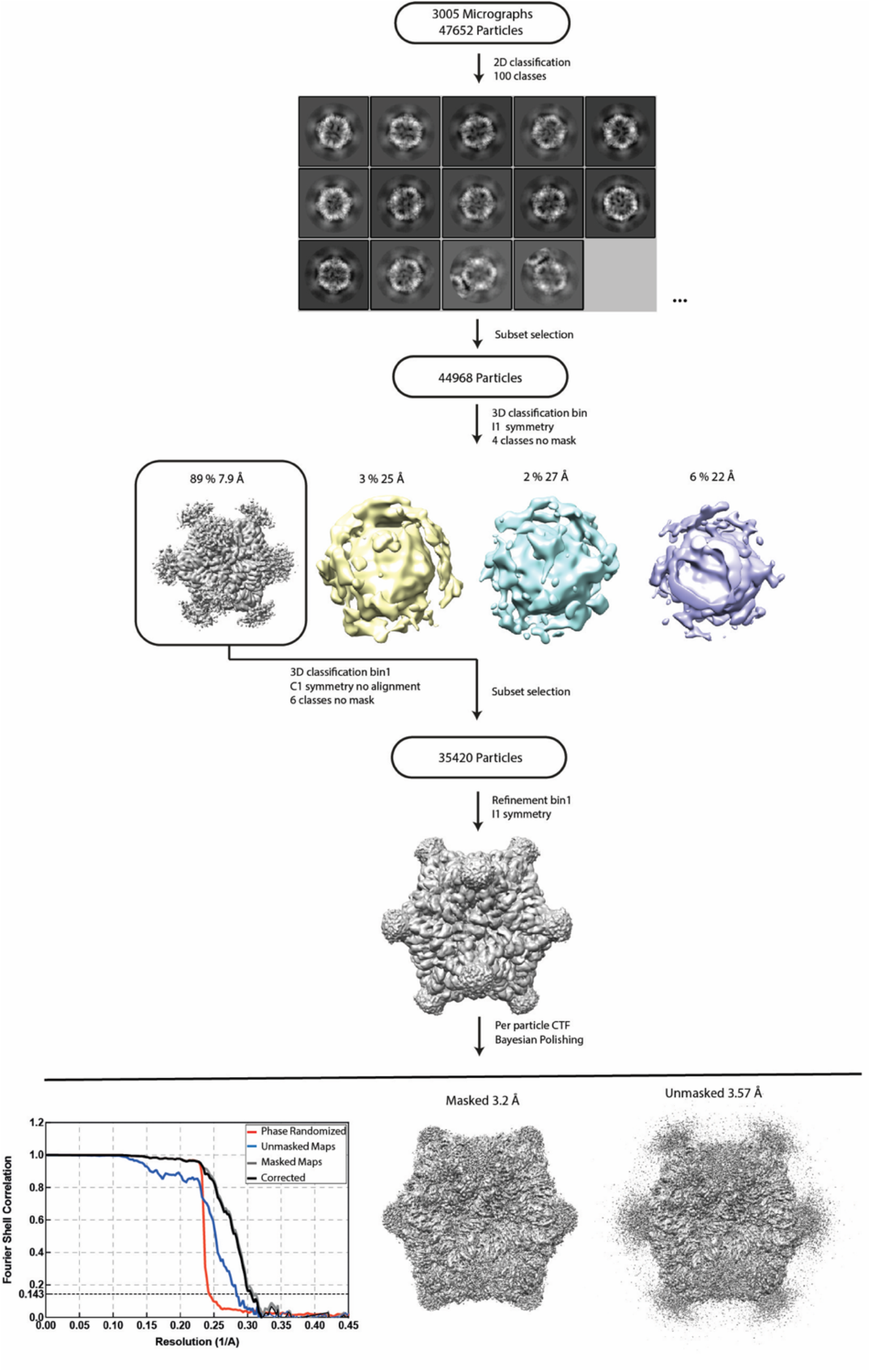
Image processing flowchart for icosahedral PNMA2 density map reconstruction. Resolution of reconstructions are determined by gold-standard Fourier shell correlation (FSC) at the 0.143 criterion.

**Supplemental Figure S6:**
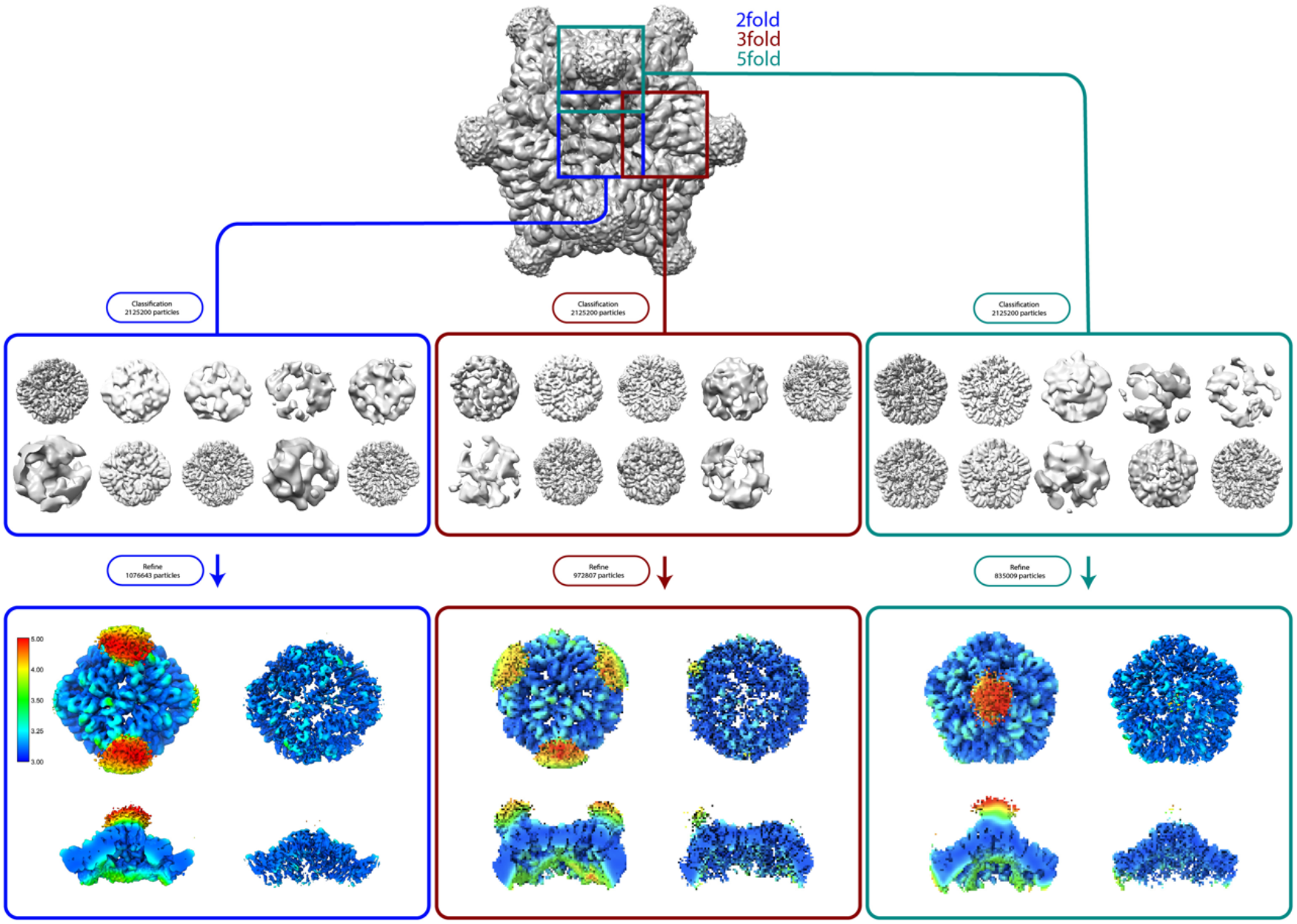
Image processing flowcharts of local density map reconstructions. For details see materials and methods. The unsharpened and sharpened maps of locally-refined five-, three- and two-fold symmetric axes are colored from high (blue) to low (red) local resolutions calculated using ResMap.

**Supplemental Figure S7:**
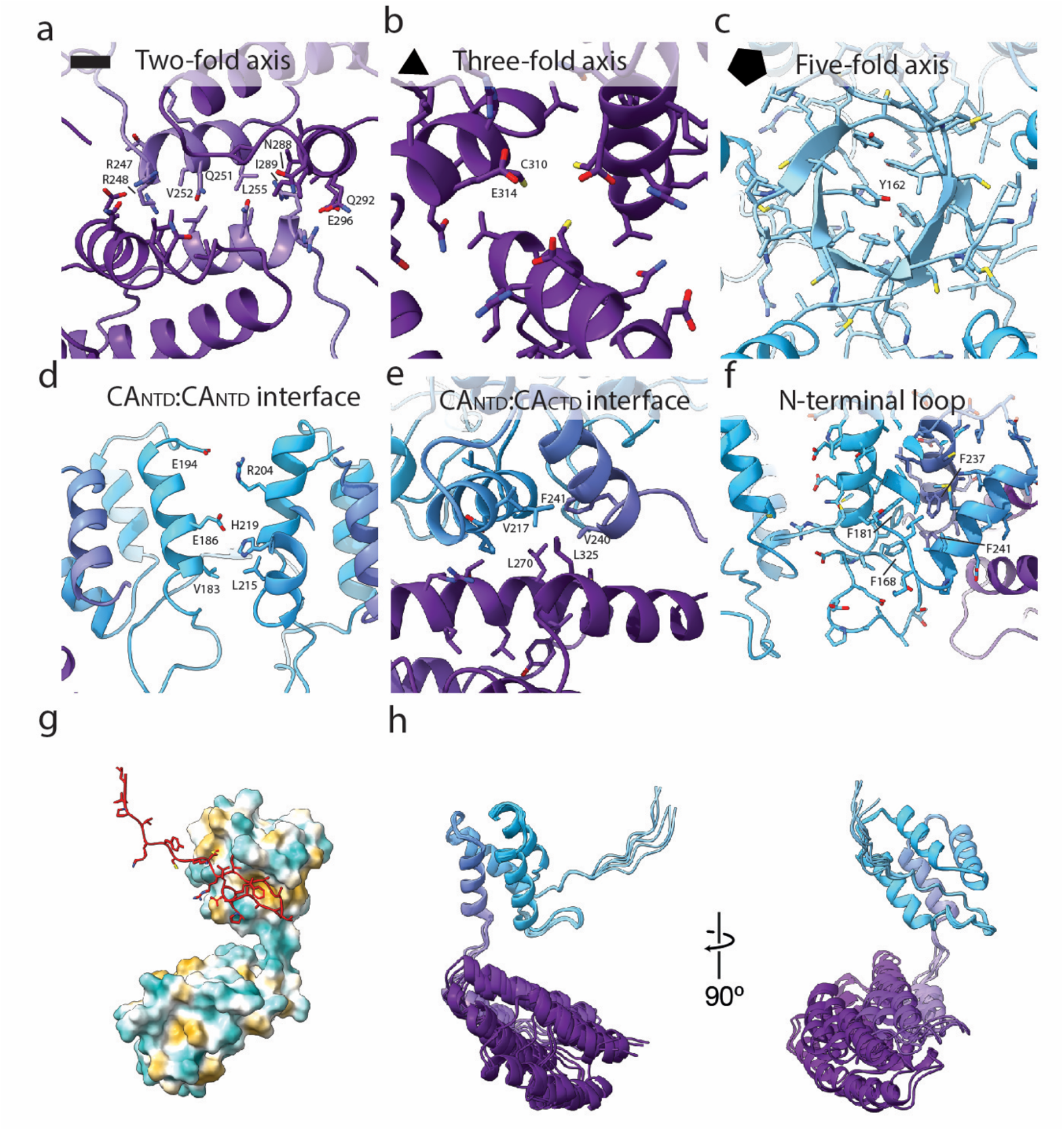
Detailed overview of mPNMA2 capsid interfaces and flexibility. **a**. The two-fold CA_CTD_ interface axis is formed between helices 5 and 7 through a hydrophobic interaction between V252 and L255, flanked by charged and neutral residues. We detect a potential salt bridge between R247 and Q292. **b**. Helix 8 constitutes the threefold CA_CTD_ interfaces, where E314 and C310 form the interface between the individual CA molecules. We don’t see any density which could be assigned to coordination of ions within this interface. **c**. The five-fold axis is constituted by residues 160-164 from each of the five CA molecules which form a short asymmetrical five-stranded beta-barrel, with one Y162 occupying the center of the capsomer. **d**. The CA_NTD_:CA_NTD_ interfaces are formed between helices 1-3. Residues in helix 1, E186 and E194, may form salt bridges with R204 and H219 of helices 2 and 3, respectively. **e**. The CA_NTD_:CA_CTD_ interface forms by docking of CACTD helix 6 residues L270 and L325 into a hydrophobic cavity formed by V217,F241 and V240 of helices 3 and 4 of the CA_NTD_. **f-g**. The N-terminal residues F168 and M165 preceding helix 1 dock into two distinct hydrophobic grooves of CA_NTD_. We note that in the rArc NMR and x-ray structure a slightly larger hydrophobic groove constitutes the binding site for e.g TARPy2 and the NMDA cytosolic tails. **h**. Local reconstructions revealed that the capsids are not perfectly icosahedral and that the relative orientation of the CA_NTD_ and CA_CTD_ domains varies (also see movie S3 and S4).

**Supplemental Figure S8:**
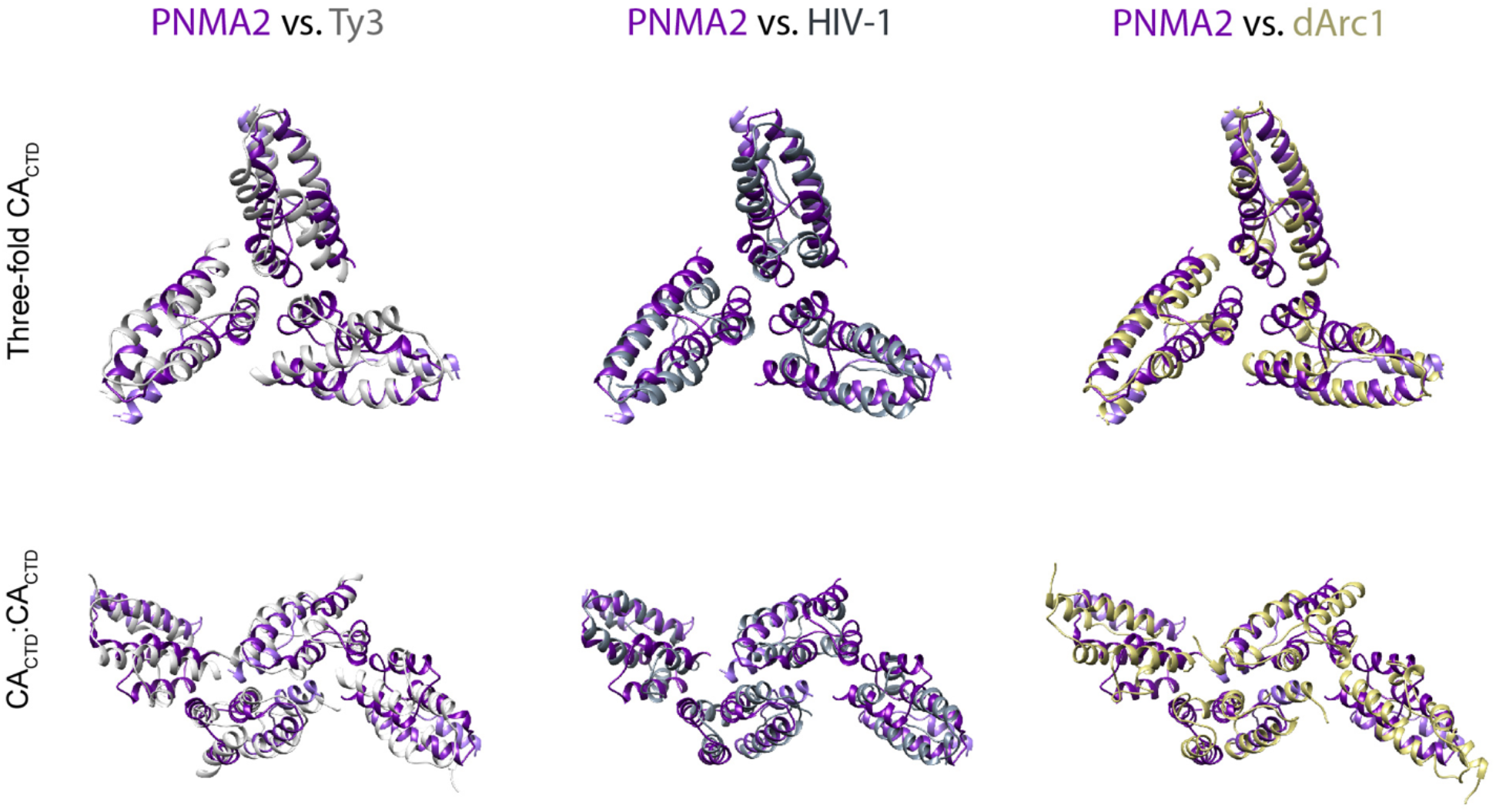
Alignment of the PNMA2 capsomer interfaces compared to dArc1, Ty3 and mature HIV capsid interfaces. The arrangement of CACTD at the five-three, and two-fold interfaces in the PNMA2 capsid is similar to the arrangement at the corresponding positions in Ty3 (light grey), mature HIV-1 (PDB ID: 5MCX; dark grey) and dArc1 (PDB ID: 6TAP; yellow), For monomer alignments see Movie S2.

**Supplemental Figure S9:**
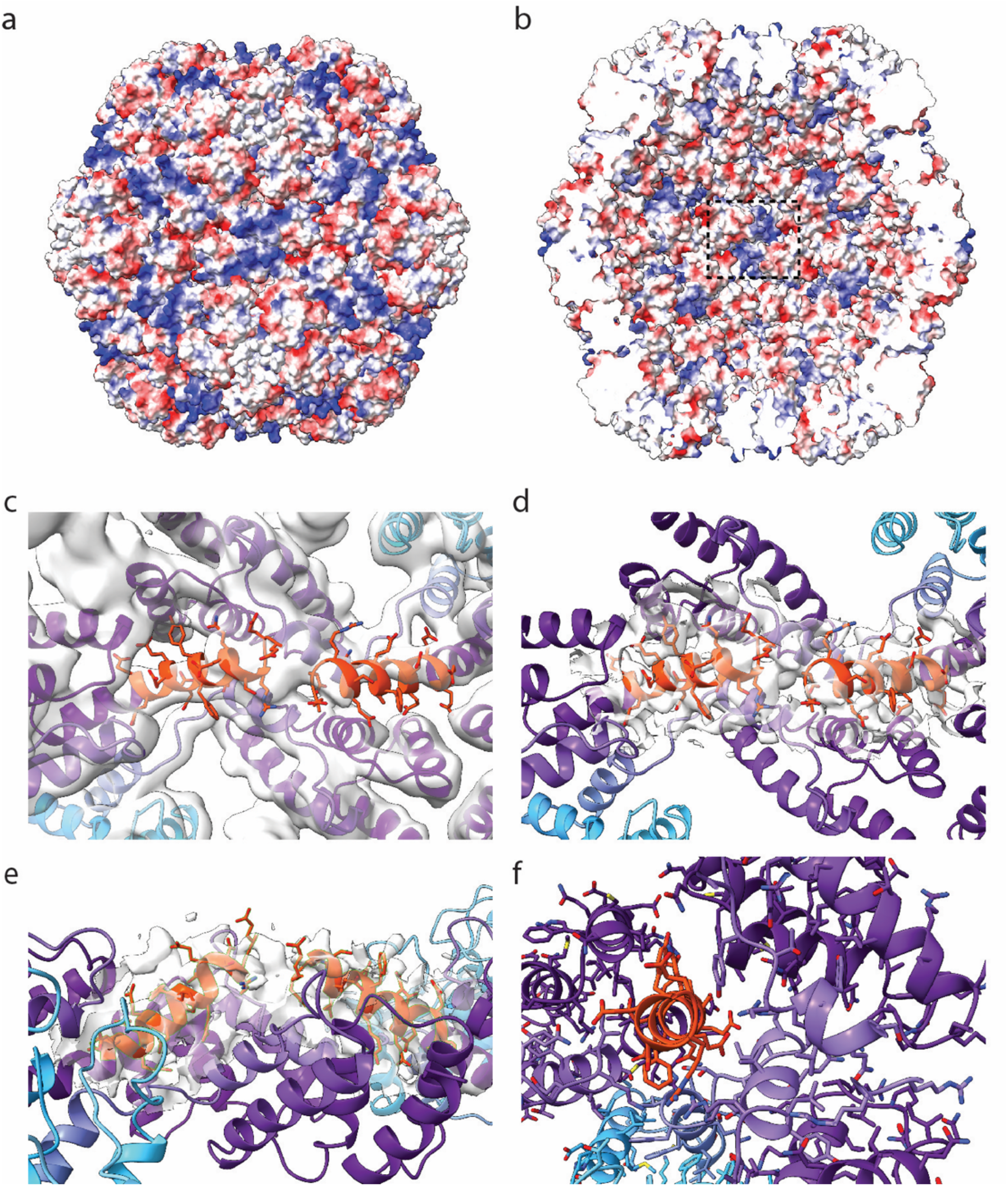
Electrostatic potential and C-terminal CA extension. **a-b**. PNMA2 capsid exterior and interior surfaces colored by electrostatic potential from -5 (red) to +5 kbT/e (blue). **c-f**. Different views of the tentative models of the C-terminal CA extension (red). The density is not well resolved in this region.

**Supplemental Figure S10.**
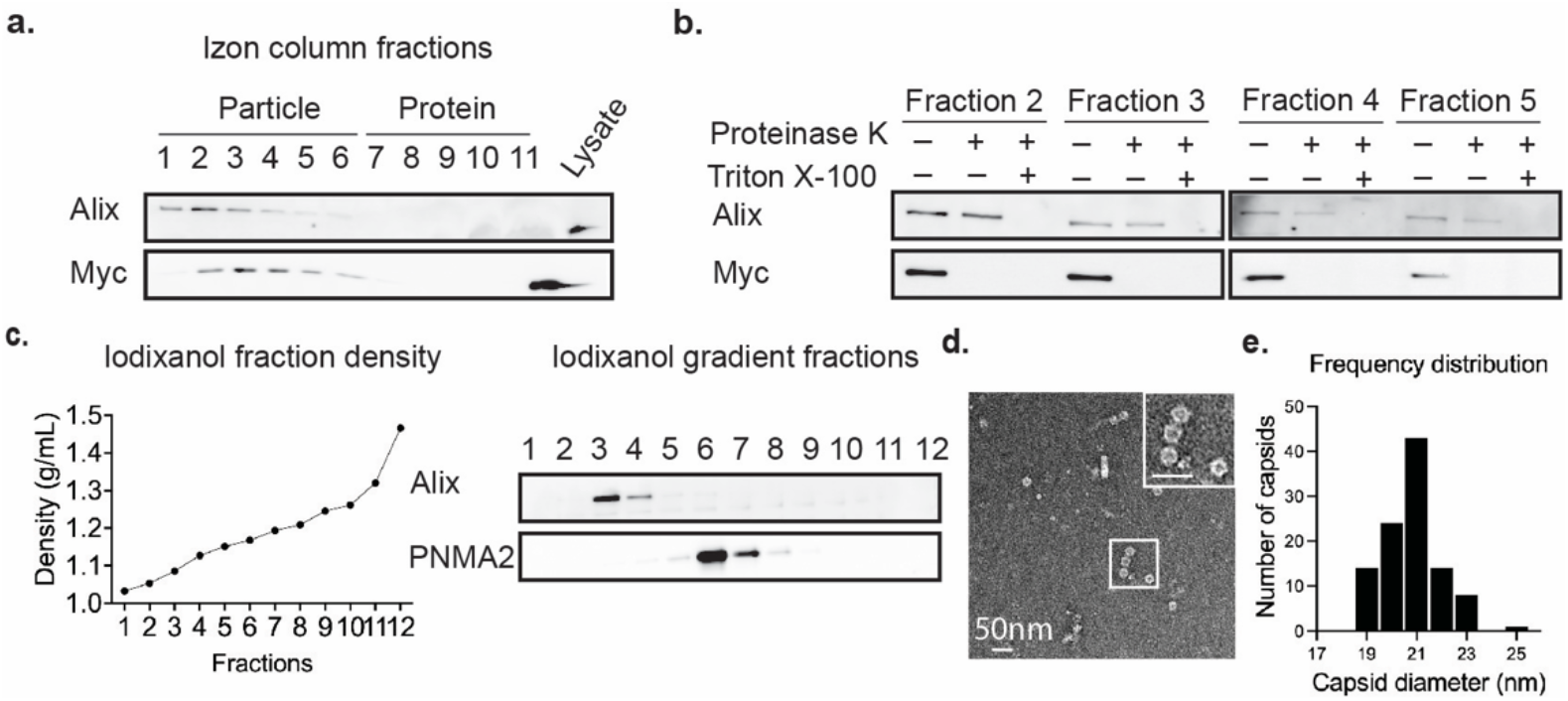
mPNMA2 is released as non-enveloped capsids by HEK 293T cells overexpressing myc-mPNMA2. **a**. Media was collected from myc-PNMA2 transfected HEK 293T cells after 24h incubation and fractionated using size exclusion chromatography (SEC). Fractions were run on a gel and blotted for Myc and ALIX. PNMA2 protein is released in early fractions that contain EV proteins. **b**. The early fractions 2-5 from SEC were blotted for Myc and ALIX. One set of fractions were incubated with Proteinase K (200µg/mL) with or without detergent present (1% Triton-X) for 10mins. Representative Western blots show that PNMA2 protein was sensitive to Proteinase K degradation without detergent present. **c**. The pooled early fractions (1-6) from SEC were fractionated using ultracentrifugation. An iodixanol gradient was used to separate proteins by density and size. Myc-PNMA2 protein was enriched in fraction 6, while ALIX was enriched in fractions 3 and 4. **d**. A representative negative-staining EM image of non-enveloped PNMA2 capsids isolated from the iodixanol gradient fraction 6 in c. **e**. The quantification of capsid diameters in iodixanol gradient fraction 6 imaged by negative-staining EM. These capsids show similar morphology and diameters with purified mPNMA2 capsids from *E*.*coli* system.

**Supplemental Figure S11.**
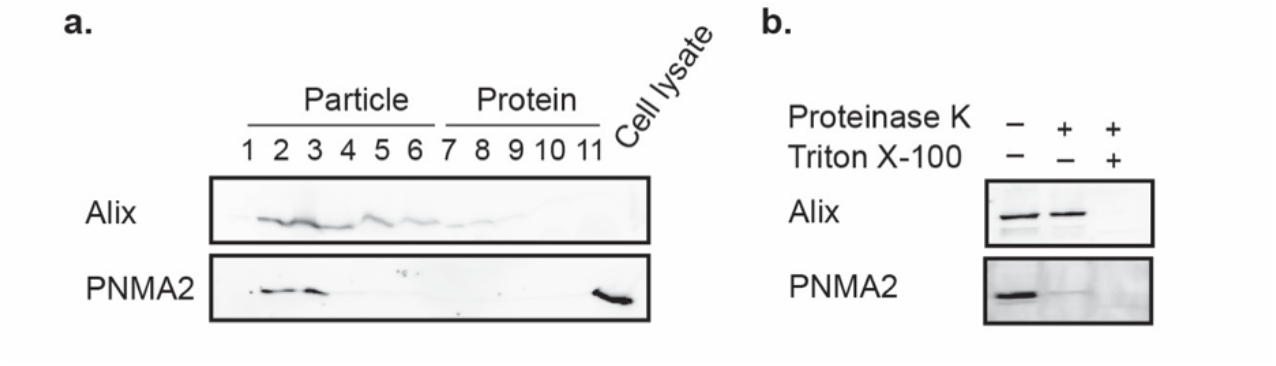
Endogenous hPNMA is released by NCI-H378 small cell lung cancer cells as non-enveloped capsids. **a**. Media was collected from NCI-H378 cell culture after 24h incubation and fractionated using size exclusion chromatography (SEC). Fractions were run on a gel and blotted for hPNMA2 protein and ALIX, a canonical EV marker. hPNMA2 protein was released in early fractions that contained EV proteins. **b**. The early fractions (1-3) from SEC were pooled and blotted for hPNMA2 and ALIX. One set of fractions was incubated with Proteinase K (200µg/mL) with or without detergent present (1% Triton-X) for 10mins. Western blots showed that hPNMA2 protein was sensitive to Proteinase K degradation without detergent present.

**Supplemental Figure S12.**
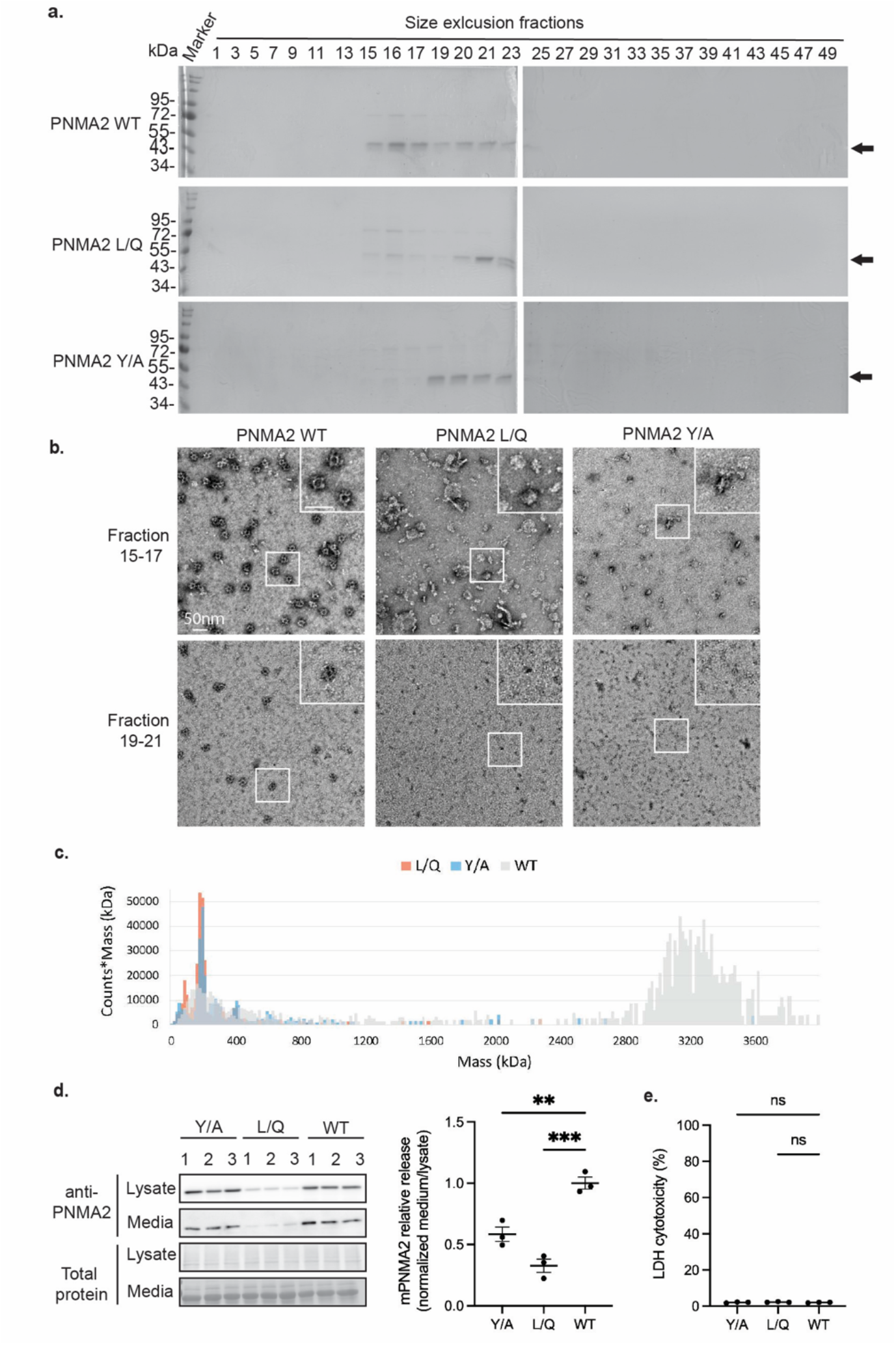
mPNMA2Y162A and L270QL325Q mutants do not form capsids and are not robustly released from cells. **a**. Representative Coomassie SDS-PAGE gels show peak fractions eluted from the size exclusion chromatography (SEC) column of purified mPNMA2, mPNMA2 L/Q, and mPNMA2 Y/A mutant proteins. mPNMA2 L/Q and Y/A proteins are shifted to later fractions. **b**. Representative negative-stained EM images of mPNMA2, mPNMA2 L/Q, and mPNMA2 Y/A proteins from pooled SEC fractions 15-17 and 19-21. mPNMA2 L/Q and mPNMA2 Y/A proteins do not form capsids. **c**. Mass distribution histograms of purified mPNMA2, L/Q, and Y/A proteins, measured by mass photometry. mPNMA2 L/Q and Y/A show a lack of high-molecular-weight assemblies (> 3000 kDa). **d**. Western blot of mPNMA2, mPNMA2 L/Q protein, and mPNMA2 Y/A expression in cell lysates and media from transfected HEK 293T cells. (***One-way ANOVA with multiple comparisons corrected by Dunnett test, P=0.0004. WT vs. Y/A: **P=0.0029; WT vs. L/Q: ***P=0.0002). **e**. LDH cytotoxicity assay to test the viability of HEK 293 cells in d. (One-way ANOVA with multiple comparisons corrected by Dunnett test, P=0.3918; WT vs. Y/A: P=0.9998; WT vs. L/Q: P=0.3853). Error bars indicate mean ± s.e.m. Abbreviation. WT: wild type; L/Q: L270QL325Q; Y/A: Y162A.

**Supplemental Figure S13.**
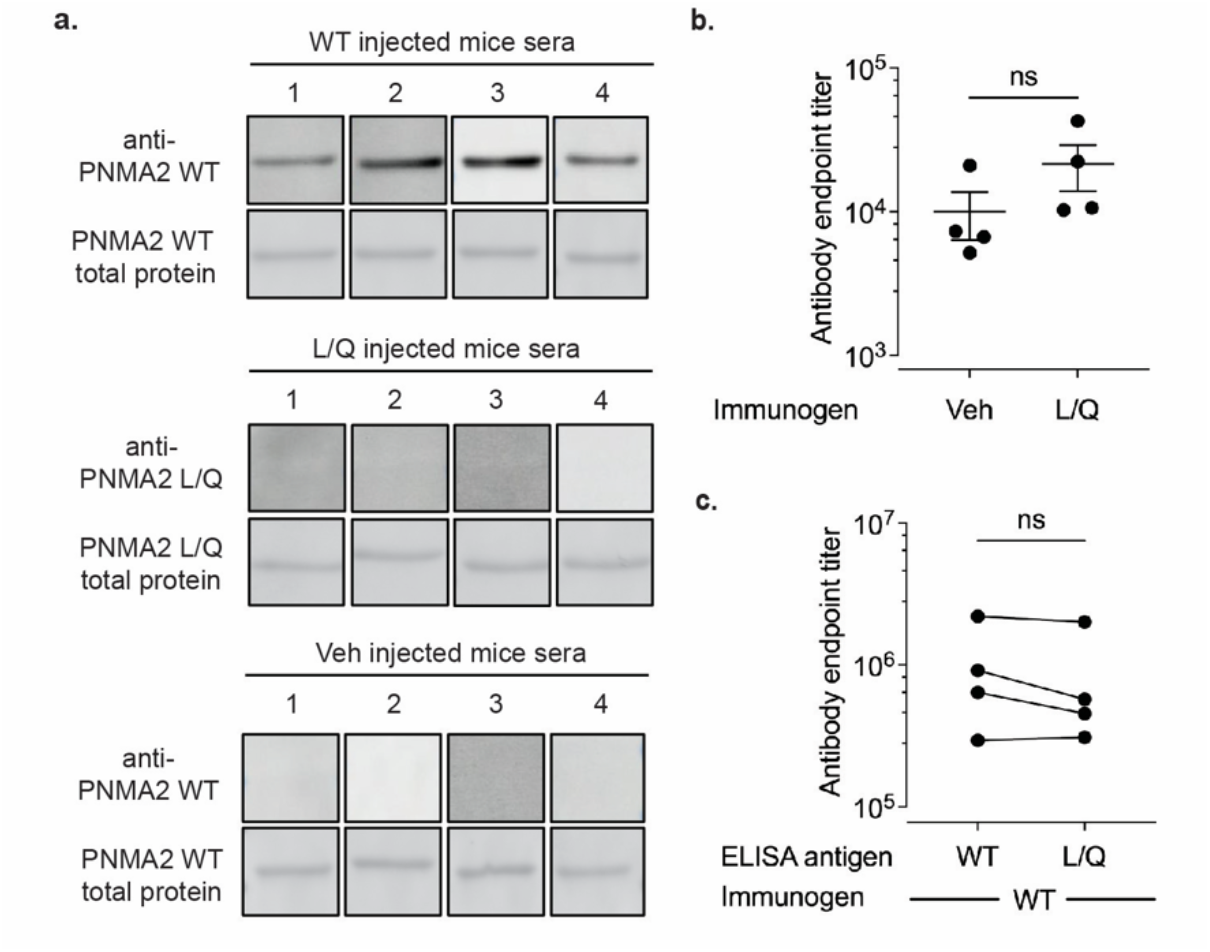
mPNMA2 capsids injected into mice induce autoantibody production. **a**. Mice were injected intraperitoneally with vehicle (n=4), 5µg purified mPNMA2 capsids (n=4) or 5µg mPNMA2 L/Q protein (n=4) and blood sera collected 3 weeks after injections. Sera collected from mice were used as the primary antibody for Western Blots of purified mPNMA2 or mPNMA2 L/Q protein. No signal was detected in sera collected from vehicle or PNMA2 L/Q protein-injected mice. **b**. Mice sera collected 3 weeks after the second injection of vehicle or 5µg mPNMA2 L/Q were analyzed for antibodies against mPNMA2 L/Q protein by ELISA. (Student t-test, P=0.2241). **c**. mPNMA2 capsids or mPNMA2 L/Q protein were used as the ELISA antigen. No difference was observed in antibody titer from mPNMA2 capsid-injected mice sera. (Paired t-test, P=0.0953). Error bars indicate mean ± s.e.m.

**Supplemental table S1.**
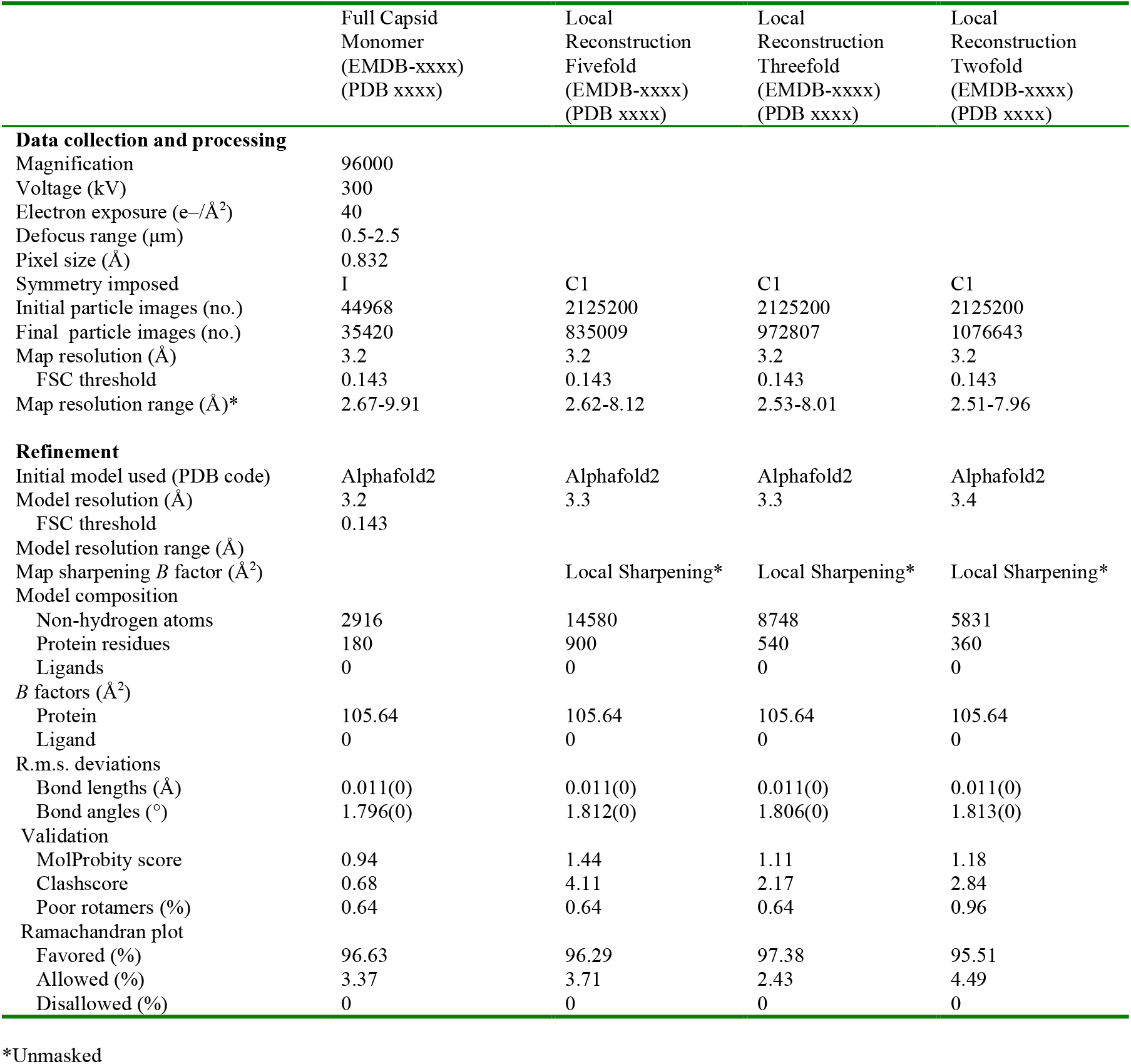
Cryo-EM data collection, refinement, and validation statistics.

